# Avant-garde assembly-line biosynthesis expands diversity of cyclic lipodepsipeptide products

**DOI:** 10.1101/560987

**Authors:** Jia Jia Zhang, Xiaoyu Tang, Tao Huan, Avena C. Ross, Bradley S. Moore

**Affiliations:** Center for Marine Biotechnology and Biomedicine, Scripps Institution of Oceanography, University of California at San Diego, La Jolla, California, USA; Department of Chemistry, University of British Columbia, Vancouver, Canada; Department of Chemistry, Queen’s University, Kingston, Ontario, Canada; Skaggs School of Pharmacy and Pharmaceutical Sciences, University of California at San Diego, La Jolla, California, USA

## Abstract

Modular nonribosomal peptide synthetase (NRPS) and polyketide synthase (PKS) enzymatic assembly lines are large and dynamic protein machines that generally undergo a linear progression of catalytic cycles via a series of enzymatic domains organized into independent modules. Here we report the heterologous reconstitution and comprehensive characterization of two hybrid NRPS-PKS assembly lines that defy many standard rules of assembly-line biosynthesis to generate a large combinatorial library of cyclic lipodepsipeptide protease inhibitors called thalassospiramides. We generate a series of precise domain-inactivating mutations in thalassospiramide assembly lines and present compelling evidence for an unprecedented biosynthetic model that invokes inter-module substrate activation and tailoring, module skipping, and pass-back chain extension, whereby the ability to pass the growing chain back to a preceding module is flexible and substrate-driven. Expanding bidirectional inter-module domain interactions could represent a viable mechanism for generating chemical diversity without increasing the size of biosynthetic assembly lines and raises new questions regarding our understanding of the structural features of multi-modular megaenzymes.

---

Nonribosomal peptide synthetase (NRPS) and polyketide synthase (PKS) enzymes are molecular-scale assembly lines that construct complex polymeric products, many of which are useful to humans as life-saving drugs. The first characterized assembly lines exhibited an elegant co-linear biosynthetic logic, whereby the linear arrangement of functional units, called modules, along an NRPS/PKS polypeptide directly correlates to the chemical structure of the product^1^. The polyketide antibiotic erythromycin^2^ and the nonribosomal peptide antibiotic daptomycin^3^ are two such examples. The core components of an assembly line elongation module include adenylation (A) or acyltransferase (AT) domains for substrate selection, thiolation (T) or carrier protein domains for covalent substrate tethering, and condensation (C) or ketosynthase (KS) domains for chain extension. Optional tailoring domains such as methyltransferase (MT), ketoreductase (KR), or dehydratase (DH) domains, if present, chemically modify building blocks or chain-extension intermediates. This one-to-one correlation between product residues and assembly line modules with the requisite catalytic domains is one feature that makes NRPS/PKS enzymes among the largest proteins found within the tree of life.

However, it is now clear that many assembly lines do not strictly abide by the rules of co-linearity. A phylogenetically distinct class of modular PKSs are *trans*-AT PKSs, which do not directly encode AT domains within modules but instead, as stand-alone enzymes, act *in trans*^4^. A separate type of NRPS, referred to as a nonlinear NRPS, deviate from the standard core domain arrangement of C-A-T and reuse a single domain more than once^5^. Several nonlinear NRPSs possess modules missing A domains that are presumably loaded by A domains from upstream modules^6–10^. Leveraging domain activities *in trans* or from different modules reduces the size of biosynthetic assembly lines and thus may represent a mechanism for minimizing modular assembly lines without sacrificing product formation.

A particularly intriguing set of hybrid NRPS-PKS assembly lines that are both nonlinear and *trans*-AT are those responsible for the biosynthesis of the thalassospiramides, a large group of immunosuppressive cyclic lipodepsipeptides^11–13^. Thalassospiramide NRPS-PKS genes have been identified in several marine *Rhodospirillaceae* bacteria and exhibit distinct architectures that range in domain and module “completeness”^13^. While it is still unclear whether all configurations are functional, thalassospiramide assembly lines representing the most “complete” and “incomplete” architectures are both capable of producing a large combination of lipodepsipeptides that vary in fatty acid and amino acid composition, order, and length^12^. Previous work^11–13^ identifying these NRPS-PKS genes and structurally characterizing their numerous and diverse chemical products led us to hypothesize that these assembly lines must operate with an unprecedented degree of nonlinearity. Furthermore, we posited that in order to generate their chemical products, thalassospiramide assembly lines must catalyze one or two rounds of pass-back chain extension, where the chain-extension intermediate is passed from a downstream module back to an upstream module through an unprecedented mechanism. Here, we report a comprehensive characterization of the thalassospiramide biosynthetic machinery from *α*-proteobacteria *Thalassospira* sp. CNJ-328 and *Tistrella mobilis* KA081020-065, which represent the most “complete” and “incomplete” assembly line architectures, respectively^13^. We present an experimentally supported and mechanistically novel biosynthetic model that invokes inter-module substrate activation and tailoring, module skipping, and pass-back chain extension, whereby the ability to pass the growing chain forward or backward is flexible and influenced by the chain length of the chemical intermediate. These newly described features accentuate the potential bidirectionality and flexibility of these large and dynamic enzymes and reveal new engineering opportunities and structural considerations for multi-modular biosynthetic enzymes.

## Results

### Cloning and heterologous expression of the *ttc* and *ttm* biosynthetic gene clusters

Close inspection of the thalassospiramide *ttc* and *ttm* gene clusters revealed that the NRPS/PKS genes from *Thalassospira* and *Tistrella*, respectively, occupy completely different genomic contexts (**Fig. 1a-b, Supplementary Table 2**). Beyond the core assembly line, composed of *ttcA*-*C* and *ttmA*-*B*, we observed no synteny between upstream or downstream genes. The *ttc* gene cluster also includes a 4’-phosphopantetheinyl transferase (PPTase), TtcD, for which there is no homolog in *ttm.* Notably absent from the genomic vicinity of both pathways are any genes encoding stand-alone AT or A domains (**Supplementary Table 2**), although both assembly lines possess a *trans*-AT PKS module and Ttm contains two NRPS modules without critical substrate-loading A domains (**Fig. 2**). While the genome of *T. mobilis* KA081020-065 is fully assembled^14^, the draft genome of *Thalassospira* sp. CNJ-328 is highly fragmented; thus, we chose to clone a conservative range for the *ttc* pathway but targeted a more limited range for *ttm* (**Fig. 1a-b**), as it would be simpler to go back and search the genome of *T. mobilis* for non-clustered, essential elements if necessary for heterologous expression.

**Fig. 1.**
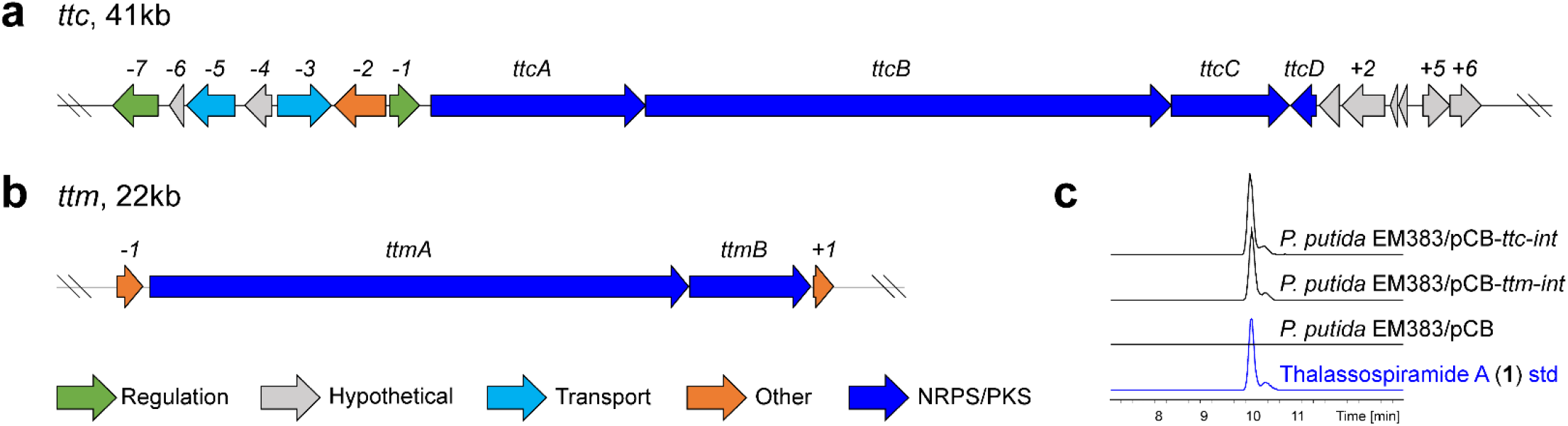
Heterologous reconstitution of thalassospiramide biosynthetic gene clusters in a *P. putida* host. Annotated genomic loci encompassing the thalassospiramide assembly line genes from (**a**) *Thalassospira* sp. CNJ-328 and (**b**) *Tistrella mobilis* KA081020-065 targeted for cloning and heterologous expression. (**c**) LC-MS analysis of extracts from an empty *P. putida* EM383 host and hosts with genomically integrated *ttc* and *ttm* pathways compared against an authentic thalassospiramide A (**1**) standard. Extracted ion chromatograms (EIC) of m/z 958.5496.

**Fig. 2.**
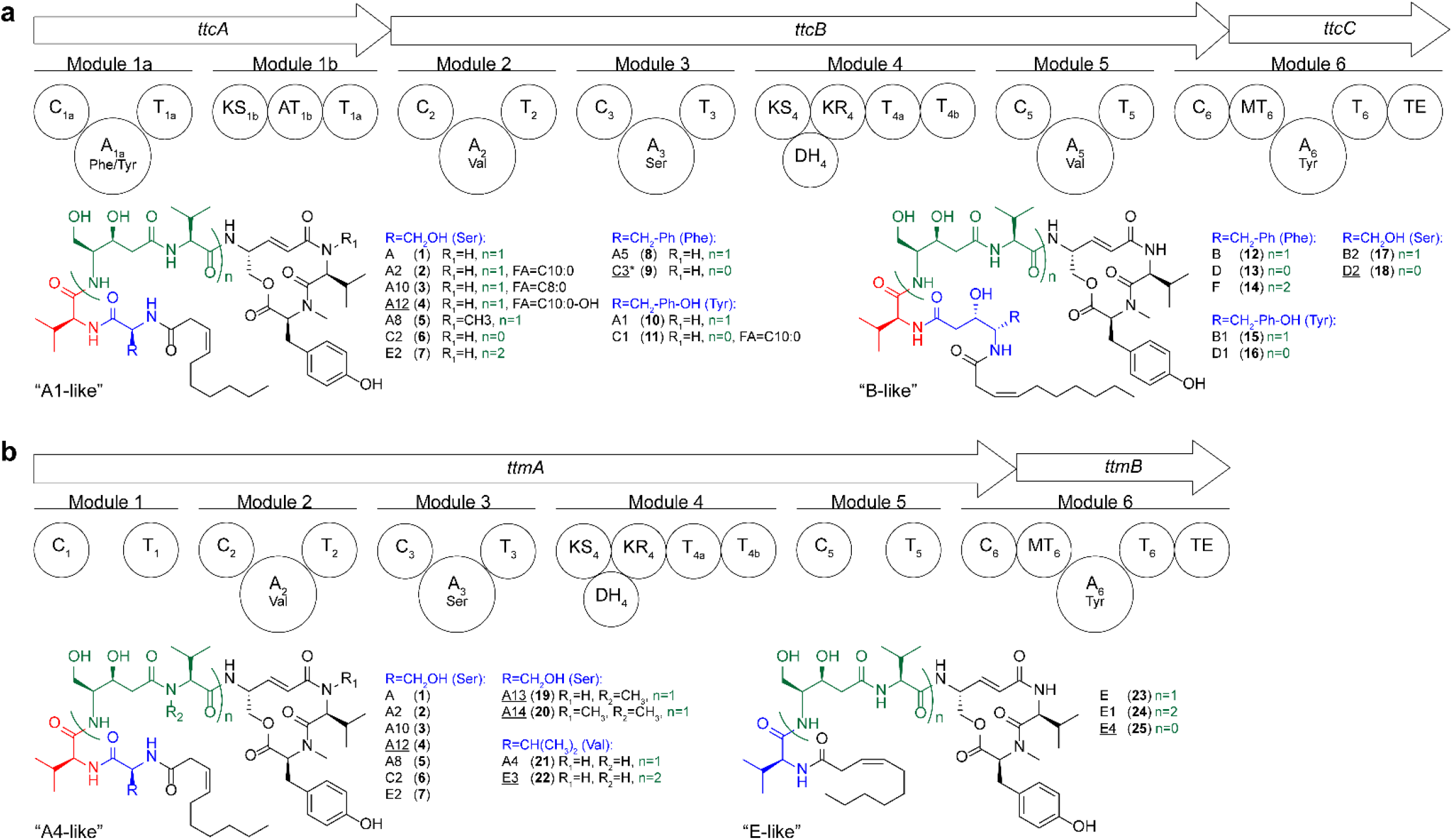
Domain and module organization of Ttc and Ttm assembly lines and structural diversity of associated cyclic lipodepsipeptide products. (**a**) Ttc assembly line and structures of a representative set of associated chemical products. (**b**) Ttm assembly line and representative associated chemical products. Analogs not previously reported are underlined. Analogs detected by LC-MS from the wild-type producer but not the heterologous host are marked with an asterisk. C, condensation; A, adenylation; T, thiolation; KS, ketosynthase; AT, acyltransferase; DH, dehydratase; KR, ketoreductase; MT, methyltransferase; TE, thioesterase.

To directly clone these pathways, a new bacterial artificial chromosome (BAC) based transformation-associated recombination (TAR) cloning vector, pCAP-BAC (pCB), was designed and constructed to enable stable maintenance of large constructs in *Escherichia coli* (**Supplementary Fig. 1**). pCB lacks host-specific integration elements, which can be introduced after cloning, to make it easier to retrofit cloned pathways with integration elements for different hosts, as we believe this is a valuable approach^15^. This also reduces the size of the vector backbone, enabling efficient generation of cluster-specific capture vectors by a simple PCR reaction using long primers with homology arms designed directly into the primer sequence. Following successful cloning of *ttc*, heterologous expression was attempted but never achieved in *E. coli*, despite efforts to perform promoter refactoring, stabilize protein expression, and co-express the pathway with various promiscuous PPTases (data not shown). Thus, we constructed a *Pseudomonas* integration cassette containing the Int-B13 site-specific recombinase^16^ and introduced it to the vector backbone to generate pCB-*ttc*-*int* (**Supplementary Fig. 1**). This construct was successfully integrated into the genome of *Pseudomonas putida* EM383^17^ (**Supplementary Fig. 2**). The same procedure was used for *ttm*, and expression of both gene clusters in *P. putida* resulted in successful production of thalassospiramide A (**1**) (**Fig. 1c**). These observations validated that the Ttc and Ttm assembly lines do not require additional pathway-specific enzymatic components, beyond what was transferred to the host and what is supplied through primary metabolism. To our knowledge, this is the first report of successful heterologous expression of a *trans*-AT pathway without co-transfer of a cognate AT. Our current hypothesis as to why heterologous expression was successful in *P. putida* but not in *E. coli* is that the *P. putida* primary metabolic AT is capable of interfacing with the *trans*-AT PKS modules, while the cognate AT from *E. coli* is not.

### Thalassospiramide structural diversity is fully recapitulated in the heterologous host

We next explored whether the heterologously expressed “complete” Ttc and “incomplete” Ttm assembly lines could produce the full suite of thalassospiramide chemical diversity. These cyclic lipodepsipeptides can be grouped into four categories based on their chemical structures and biosynthetic origin^11–13^. A subset of representative structures is shown in **Fig. 2**. Products of Ttc can incorporate serine, phenylalanine, or tyrosine as the first amino acid residue (colored blue in **Fig. 2**), which can be C2 extended to generate “B-like” as opposed to “A1-like” thalassospiramides. Alternatively, Ttm incorporates serine or valine in the first position, then includes or omits a linker valine residue (colored red in **Fig. 2**) to generate “A4-like” or “E-like” thalassospiramides, respectively. Both assembly lines produce “A-like” thalassospiramides at greater relative abundance than their “B-like” or “E-like” counterparts.

Further elements that contribute to structural diversity include the N-terminal fatty acid, which is predominantly C10:1(Δ3) but can also be fully saturated C10, C8, and C10-hydroxy fatty acids (**Fig. 2**, **Supplementary Fig. 10**), and the pattern of N-methylation, which is predominantly limited to the final tyrosine residue of the cyclic peptide core but can also extend to the adjacent valine residue for Ttc and further to linker valine residues for Ttm (**Fig. 2**, **Supplementary Fig. 11-12**). Finally, the number of ser-C2-val linker units (colored green in **Fig. 2**) can be 0, 1, or 2 for both assembly lines, which presumably arises from passage of chain extension intermediates from module 4 back to module 2 or module 5 back to module 3. Based on the different possible combinations of these variables, each assembly line can theoretically generate well over 100 compound analogs.

LC-MS-MS analysis revealed that nearly all analogs detected from the wild-type strains are also produced by the heterologous host, with some subtle differences in relative production levels. Most significantly, the host has a greater propensity to incorporate non-standard fatty acids (not C10:1) than wild-type producers (**Supplementary Fig. 3**). Overall titers of most analogs are comparable between wild-type and host and in some cases greater in the host (**Supplementary Fig. 3**).

### Characterization of non-assembly line genes

To determine whether co-transferred genes beyond the core assembly line affect thalassospiramide titer or product distribution, we performed targeted deletions of all non-assembly line genes within *ttc*. Previous work using the broad-host-range expression vector, pCAP05^15^, demonstrated that no upstream genes (*-7* through *-1*) are essential, although heterologous expression using this vector system, which exists as a self-replicating plasmid in the host, produced low yields and proved to be genetically unstable over time (**Supplementary Fig. 4a**). Targeted deletion of *ttc −1*, +*1*, +*2*, +*3*, and +*4* in the stable pCB-*ttc*-*int* expression construct had no impact on thalassospiramide production; however, deletion of the putative PPTase TtcD resulted in an approximately five-fold reduction in levels of thalassospiramide A (**1**), which was restored upon genetic complementation of *ttcD* via Tn7 transposition^18^ (**Supplementary Fig. 4-5**). This observation suggests that the single native *P. putida* PPTase is capable of activating Ttc carrier proteins, albeit not as effectively as TtcD. The *P. putida* PPTase is clearly capable of activating Ttm carrier proteins, as no cognate PPTase is present in the *ttm* gene cluster and thalassospiramides are still produced in the host. However, co-expression of TtcD with the Ttm assembly line resulted in an approximately two-fold increase in levels of thalassospiramide A (**1**) (**Supplementary Fig. 4-5**). Quantitative analysis of the impact of TtcD on production of all thalassospiramide analogs suggests that the PPTase favorably biases production of analogs that incorporate one or more ser-C2-val linker units and thus may promote pass-back chain extension through an unknown mechanism (**Supplementary Fig. 5**).

### Inactivation and testing of assembly line domains

Taken together, the results of the heterologous expression and gene deletion experiments strongly suggest that thalassospiramide structural diversity is generated directly from the multi-modular assembly line itself and does not involve accessory enzymes or tailoring domains beyond what is provided from primary metabolism, which is both remarkable and somewhat unexpected. Although certain modules appear to be missing domains based on retro-biosynthetic analysis of thalassospiramide chemical structures, such as KR_1b_, AT_4_, and MT_5_ from Ttc and A_1_, MT_2_, AT_4_, MT_5_, and A_5_ from Ttm, all necessary core and tailoring domains are present somewhere along the assembly line, including a DH domain in module 4 that was not previously annotated (**Supplementary Fig. 6, Fig. 2**). Thus, we set out to investigate the predicted inter-module activity of assembly line domains.

Our initial approach leveraged gene deletion and complementation tools, focusing first on the smaller *ttcC* that encodes the four-domain terminal module 6. Despite it harboring the lone assembly-line MT domain, some thalassospiramides, such as A8 (**5**), are unusual in containing a ‘misplaced’ penultimate N-methylated valine residue in addition to the conserved terminal N-methylated tyrosine. To explore whether MT_6_ can methylate both A5 and A6 substrates, we deleted *ttcC*, resulting in complete loss of thalassospiramide production. Complementation with wild-type TtcC restored thalassospiramide production, while complementation with the mutant TtcC-G234D, in which MT_6_ has been selectively inactivated^19^, resulted in dramatic reduction and complete loss of thalassospiramide A (**1**) and A8 (**5**), respectively, and formation of a new product with MS1 and MS2 spectra consistent with desmethyl thalassospiramide A, or thalassospiramide A15 (**28**) (**Supplementary Fig. 7, Supplementary Fig. 13**). This result confirms our hypothesis that MT_6_ can act within the upstream module 5 of TtcB. Furthermore, it suggests that pass-back chain extension occurs between modules 4 and 2 as opposed to 5 and 3, as promiscuous methylation is confined to the cyclic valine residue for Ttc. As module 5 of Ttm does not possess an A domain and presumably borrows the activity of A_2_, this could explain why promiscuous methylation can extend to upstream valine residues for Ttm but not for Ttc.

We attempted to use the same approach to characterize other thalassospiramide domains; however, efforts to perform PCR mutagenesis in *ttcA*, which is over 6.5 kb, and *ttcB* and *ttmA*, which are both over 15.5 kb, together with challenges in Tn7 transposition-based complementation, were ultimately unsuccessful. Therefore, we established new methodology combining oligo recombineering with CRISPR-Cas9 counter selection to introduce facile and precise point mutations to large DNA constructs cloned into the pCB vector backbone. Previous work demonstrated that oligo recombineering combined with CRISPR-Cas9 counter selection could greatly enhance the rate of precise and marker-less genome editing in *E. coli*, both by selecting against unedited cells as well as inducing recombination triggered by generation of dsDNA breaks^20^. Thus, we developed a similar system for editing cloned DNA constructs instead of genomic DNA and, using this method, selectively inactivated a series of assembly line domains to directly interrogate their role in thalassospiramide biosynthesis.

Consistent with our annotation of C_1a_ of TtcA as a starter C domain responsible for fatty acylation of the first amino acid residue^21^, C_1a_ inactivation completely abolished production of all thalassospiramides (**Fig. 3a**). We did not detect any masses corresponding to core peptides lacking an N-terminal fatty acid, suggesting that the assembly line can only generate lipopeptide products. In contrast, inactivation of A_1a_ using two different point mutations, TtcA-G631D and TtcA-K972A, resulted in essentially complete loss of all thalassospiramides incorporating phenylalanine or tyrosine as the first residue but maintained or enhanced production of analogs incorporating serine in the first position (**Fig. 3a**). Control inactivation of A_3_ using the analogous lysine to alanine mutation (TtcB-K2045A) resulted in complete loss of all thalassospiramide production, as did selective inactivation of T_1a_ (**Fig. 3a**). These results suggest that A_3_ directly adenylates T_1a_ during biosynthesis of analogs that incorporate serine as the first amino acid residue, which is in fact preferred, as thalassospiramides incorporating serine in the first residue predominate.

**Fig. 3.**
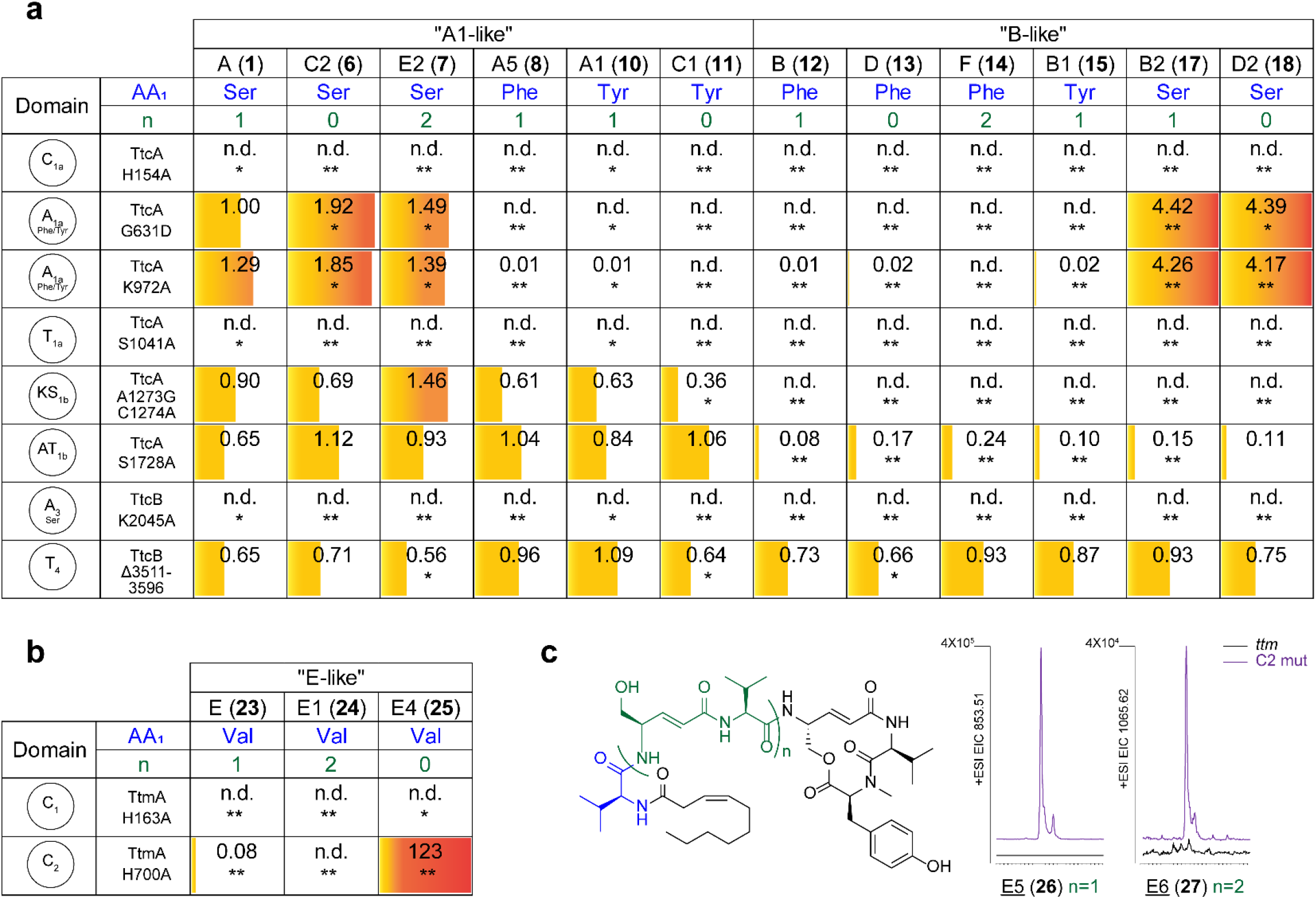
Selective inactivation of assembly line enzymatic domains alters product formation. Changes in production level of thalassospiramide analogs from (**a**) Ttc and (**b**) Ttm assembly lines upon selective inactivation of specific enzymatic domains. Domain and precise amino acid mutations are listed in the first two columns. Fold-change in MS ion intensity is indicated by number, color intensity, and fill proportion; half-filled boxes indicate no change compared to wild-type, while empty boxes indicate analogs were not detected (n.d.) from the mutant. Statistical significance was calculated using a two-tailed Student’s t-test; n=3, *p<0.05, **p<0.005. (**c**) Structures and EICs of new thalassospiramide analogs produced upon TtmA C_2_ inactivation; full MS1 and MS2 spectra provided in **Supplementary Fig. 19** and **20**.

We hypothesized that “B-like” thalassospiramides from Ttc arise through PKS module 1b extension and, correspondingly, that “A1-like” thalassospiramides arise through module 1b skipping. Consistent with that hypothesis, inactivation of KS_1b_ resulted in complete loss of “B-like” analogs but maintained production of “A1- like” analogs, although some yields were slightly reduced (**Fig. 3a**). Surprisingly, although AT_1b_ inactivation reduced production of “B-like” analogs, almost all could still be detected by LC-MS analysis at ~8-24% of wild-type production levels (**Fig. 3a**). As the AT_1b_ mutation is expected to abolish enzymatic activity, we propose that the module 4-interacting *trans*-AT can partially complement AT_1b_ activity.

We previously hypothesized that the tandem T domains in module 4 might be important for determining whether chain extension intermediates are passed forward or backward^12^, although literature precedents indicated that multiple T domains do not change product identity but instead increase flux or yield^22^. We attempted to use the same oligo recombineering/CRISPR-Cas9 method to selectively inactivate T_4a_ and T_4b_. However, CRISPR-Cas9 targeting did not result in oligo incorporation but instead recombination across the two very similar T domain sequences to generate an in-frame deletion of residues 3511 to 3596 in TtcB. The resultant mutant protein contained only a single chimeric T domain composed of 46% of the N-terminus of T_4a_ and 54% of the C-terminus of T_4b_ (**Supplementary Fig. 9**). Transfer of this construct to the heterologous host revealed that all thalassospiramide analogs could still be produced, including those incorporating linker units arising from pass-back chain extension, albeit in decreased yields (**Fig. 3a**). This result proves that tandem T domains are not essential for bi-directional chain passage. Moreover, production of all analogs was affected equally, suggesting that the tandem T domains do not influence product identity.

Finally, we predicted that “E-like” thalassospiramides produced by Ttm arise from module 2 skipping, analogous to module 1b skipping in Ttc. Thus, if pass-back chain extension occurs through modules 5 and 3, C_2_ inactivation in Ttm would preserve production of all “E-like” analogs, as C_2_ is completely skipped in this model. While C_1_ inactivation abolished production of all thalassospiramides, analogous to C_1a_ inactivation in Ttc, C_2_ inactivation reduced (but did not abolish) production of thalassospiramide E (**23**) to ∼8% of wild-type levels (**Fig. 3b**). This result confirms our hypothesis that most pass-back events occur between modules 4 and 2. Furthermore, we observed a more than 100-fold increase in levels of thalassospiramide E4 (**25**), suggesting that the inability to pass growing chains back via C_2_ forces the assembly line to pass intermediates forward, resulting in enhanced production of the “premature” termination product E4 (**25**) (**Fig. 3b**). However, we also observed the formation of two new compounds not previously detected with MS1 and MS2 spectra consistent with the structures of thalassospiramides E5 (**26**) and E6 (**27**), indicating that pass-back between modules 5 and 3 can occur but correlates with additional substrate dehydration (**Fig. 3c, Supplementary Fig. 18-19**).

### Avant-garde features of thalassospiramide biosynthesis

The results of Ttm C_2_ inactivation provide a clear mechanistic insight into the flexibility and control of pass-back chain extension during thalassospiramide biosynthesis, as illustrated in **Fig. 4**. Selective inactivation of C_2_ does not affect early stages of “E-like” thalassospiramide biosynthesis, during which A_2_ loads T_1_ with valine and then module 2 is skipped following appendage of the N-terminal fatty acid. Assembly then proceeds through modules 3 and 4, but the chain-extension intermediate does not undergo immediate dehydration and may not be directly accessible to DH_4_. At this point, the assembly line would normally pass the chain extension intermediate from module 4 back to module 2 via C_2_, which is favored based on normal product distribution, as E4 (**25**) is normally produced at very low abundance. We propose the assembly line “measures” chain length within PKS module 4 and adopts a conformation that promotes passage backward to module 2 instead of forward to module 5. However, C_2_ inactivation forces the chain-extension intermediate forward, making it accessible to DH_4_ before it enters module 5. Once within module 5, the intermediate would normally proceed directly to module 6, resulting in formation of thalassospiramide E4 (**25**). Consistent with this proposal, C_2_ inactivation drives a substantial increase in levels of E4 (**25**) compared to wild-type. However, chain length can also be “measured” within NRPS module 5, prompting the assembly line to catalyze pass-back of a subset of intermediates that have already undergone dehydration from module 5 back to module 3, resulting in eventual formation of new products E5 (**26**) and E6 (**27**). This result provides additional evidence that the substrate becomes accessible to DH_4_ just as it is passed forward to module 5. Furthermore, it suggests that biosynthesis is flexible and controlled by a mechanism that measures chain length within multiple modules. If we assume that dehydration of the linker unit is a signature for module 5 progression, there is evidence that pass-back between modules 5 and 3 occurs at low frequency under normal conditions, perhaps as an additional checkpoint, as we can identify thalassospiramides E7 (**29**), E8 (**30**), E9a (**31**) and E9b (**32**) by LC-MS (**Supplementary Fig. 20-23**), which are all produced by wild-type Ttm and have undergone additional rounds of “premature” linker dehydration.

**Fig. 4.**
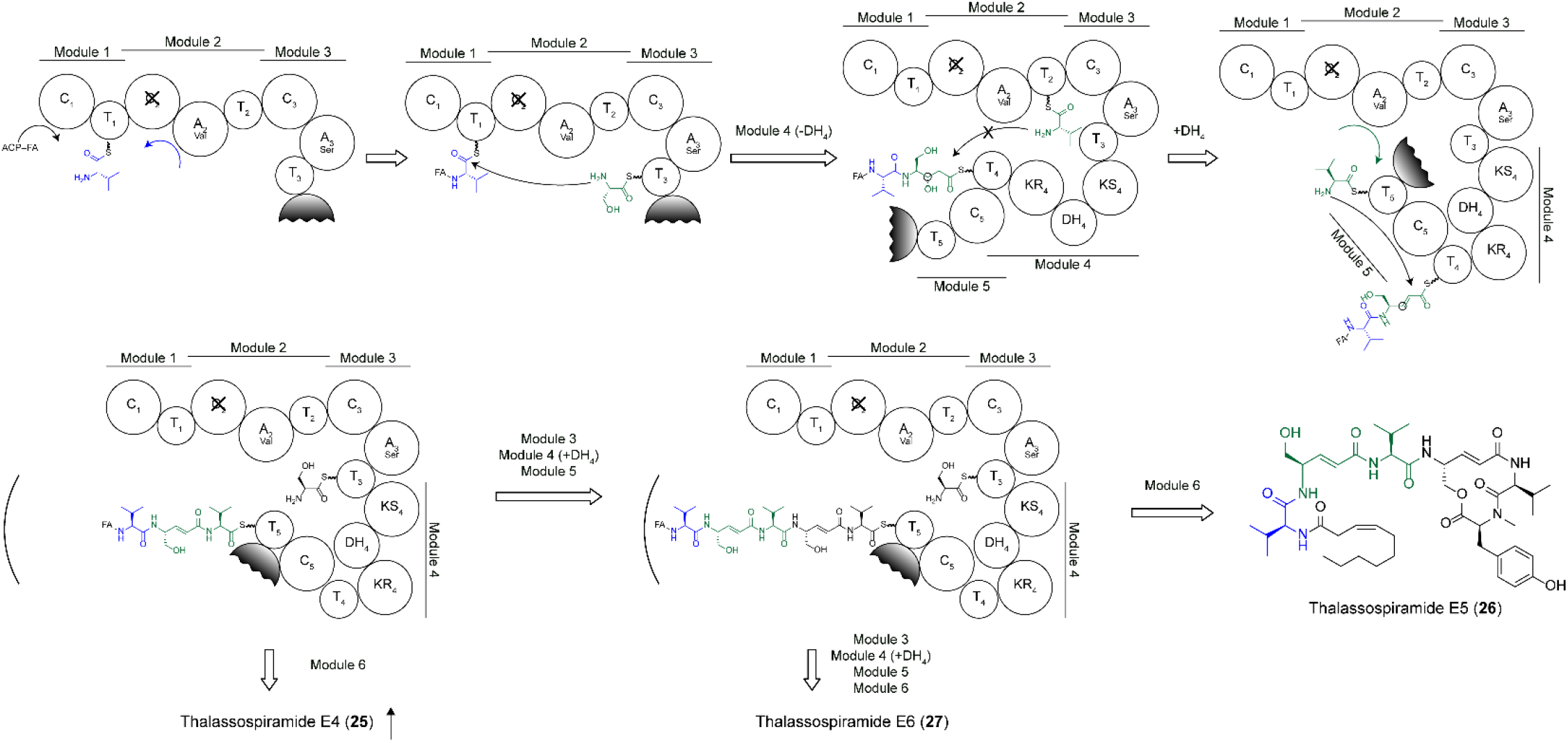
Model for thalassospiramide biosynthesis by Ttm C_2_ inactivation mutant. Valine is loaded onto T_1_ by A_2_ and C_1_ catalyzes addition of an activated fatty acid (FA) bound to an acyl carrier protein (ACP) or coenzyme A. Module 2 is skipped, and the fatty valine is extended directly onto T_3_ via C_3_. Chain extension proceeds normally to module 4, where the substrate is not immediately dehydrated and would normally be passed back to module 2 via C_2_, perhaps based on chain length. However, C_2_ inactivation forces the intermediate forward, where it becomes transiently accessible to DH_4_ and is dehydrated before extension onto T_5_, which is loaded with valine by A_2_. Direct progression through module 6 results in formation of thalassospiramide E4 (**25**), which is substantially increased as a result of C_2_ inactivation. Alternatively, passage from module 5 back to module 3 results in generation of new thalassospiramide analogs E5 (**26**) and E6 (**27**), which undergo one or two additional rounds of chain extension, respectively, through modules 3-5.

We can also propose a full mechanistic model for thalassospiramide A (**1**) biosynthesis via the Ttc assembly line (**Fig. 5**). T_1a_ is preferentially adenylated with serine through the downstream A_3_ domain. To our knowledge, this is the first report of an A domain activating a carrier protein within an upstream module that already possesses its own active A domain^12^. Subsequently, module 1b is skipped during the formation of “A1- like” thalassospiramides. Normal chain extension proceeds from modules 2 to 4, at which point we hypothesize that the substrate is sequestered from DH_4_ activity and module 5 entry based on chain length. We propose that during the formation of “B-like” thalassospiramides, ketoreduction to generate the upstream statine-like amino acid residue occurs at this time, analogous to what occurs during bacillaene biosynthesis^23^. Alternatively, as evidenced by the ability of the *trans*-AT acting within module 4 to complement inactivated AT_1b_, it is also possible that KR_4_ can act *in trans* on substrates within module 1b. The chain extension intermediate within module 4 is then passed back to module 2, where it undergoes another round of linear chain extension through modules 2 to 4 (**Fig. 5**). Now, the intermediate has reached “sufficient” chain length and becomes accessible to DH_4_ before transfer to module 5. MT_6_ promiscuously methylates valine residues activated by module 5, perhaps due to its unique positioning at the N-terminus of A_6_, to generate analogs such as thalassospiramide A8 (**5**). Finally, normal progression through modules 5 and 6 results in the formation of thalassospiramide A (**1**), the most abundant product of both Ttc and Ttm.

**Fig. 5.**
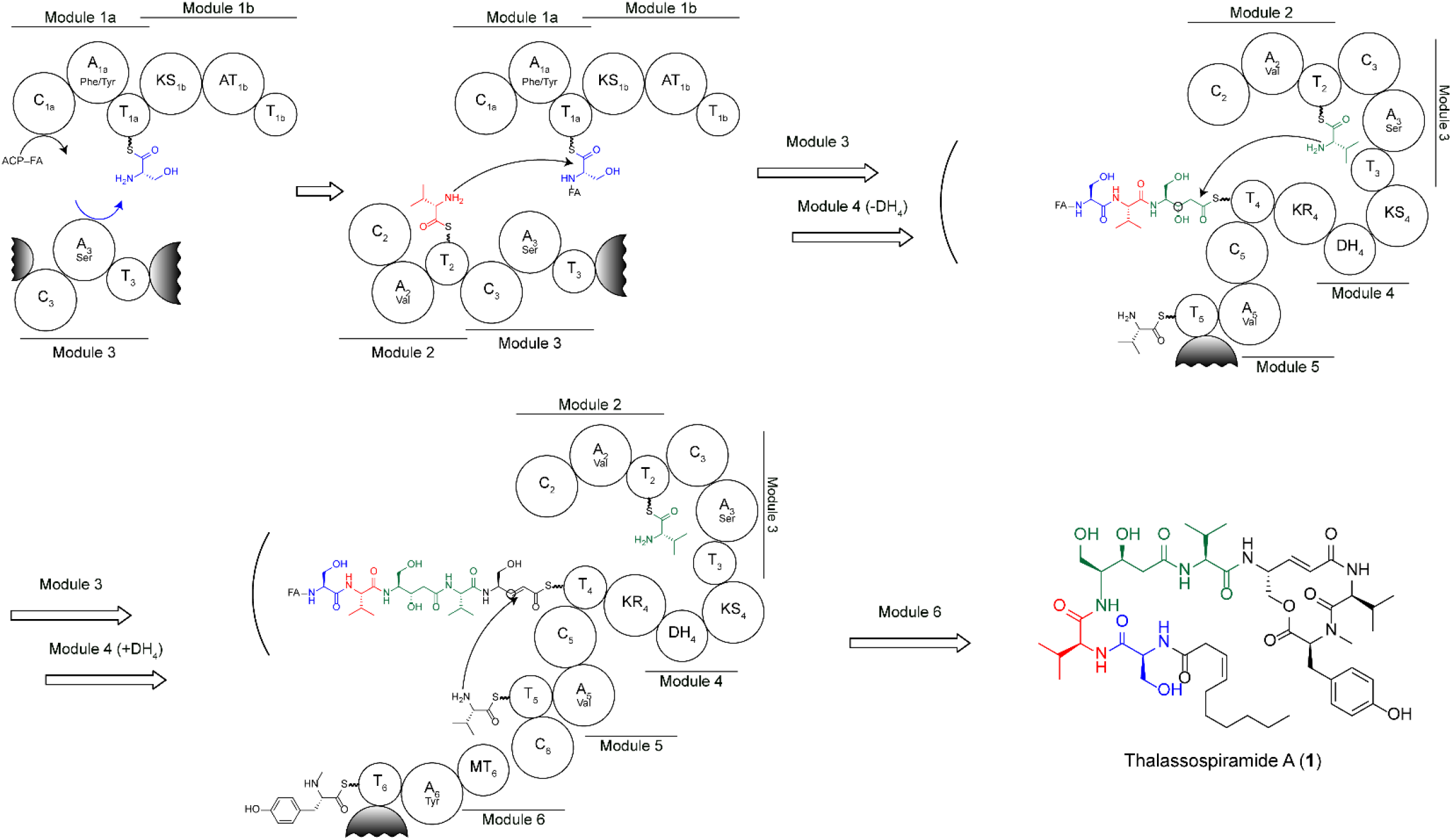
Model for thalassospiramide A biosynthesis by Ttc. Serine is loaded onto T_1a_ by A_3_ and C_1a_ catalyzes addition of an activated FA bound to an ACP or coenzyme A. Module 1b is skipped, and the fatty serine is passed directly to T_2_ via C_2_. Chain extension proceeds normally to module 4, where the substrate is sequestered from DH_4_, perhaps within an enzyme binding pocket. The intermediate is passed from module 4 back to module 2 via C_2_, where it undergoes another round of chain extension through modules 3 and 4. Upon return to module 4, the intermediate is now acted upon by DH_4_, perhaps due to a conformational change associated with longer chain length, and extended forward through modules 5 and 6 to generate thalassospiramide A (**1**).

## Discussion

Multi-modular assembly lines are large and dynamic enzymes that are nonetheless generally considered to be rather inflexible^24^. In this work, we demonstrate their potential elasticity and ability to catalyze bidirectional and nonlinear passage of chain extension intermediates to generate chemical diversity.

Several new features of multi-modular NRPS/PKS biosynthesis are described in this work. Thalassospiramide assembly lines catalyze several instances of inter-module substrate activation and tailoring. While A domain supplementation has been previously reported^6–10^, prior examples have been limited to upstream domains supplementing downstream modules, often encoded on separate proteins, and only within modules that lack their own A domains. For Ttc, A_3_ adenylates T_1a_ with serine more frequently than A_1a_ does with phenylalaine or tyrosine, although A_1a_ is active and participates in biosynthesis of numerous thalassospiramide analogs. For Ttm, both A_2_ and A_3_ deliver substrates to T_1_, but only A_3_ adenylates T_5_. Furthermore, only A_2_ loads T_1_ if module 2 is skipped, as all “E-like” analogs incorporate valine as the first amino acid, perhaps due to the specificity of C_3_^25^. MT_6_ promiscuously methylates valine residues activated by A_5_ of Ttc and A_2_ of Ttm. Finally, the *trans*-AT interacting with module 4 can supplement the upstream PKS module 1b of Ttc to a certain extent. Our model does not take possible homodimerization through PKS modules into consideration, as we do not know how the oligomeric state of hybrid NRPS-PKS enzymes influences thalassospiramide biosynthesis.

Thalassospiramide assembly lines also catalyze programmed module skipping. Forward module skipping has been previously observed in PKS^26–28^ and NRPS^29–31^ systems, both naturally and as a byproduct of assembly line engineering. Ttc and Ttm catalyze analogous skipping of PKS module 1b and NRPS module 2, respectively, although NRPS module 2 does not sit at an enzyme junction but is embedded within a very large polypeptide. Perhaps as a result, module skipping is favored in Ttc but disfavored in Ttm based on product distribution.

Finally, thalassospiramide assembly lines catalyze pass-back chain extension. To our knowledge, this has not been previously reported in modular assembly line systems, although it is analogous to “iteration” observed in the fungal beauvericin and bassianolide synthetases^32^, where intermediates are passed back and forth between adjacent modules. We argue that pass-back chain extension is distinct from iteration interpreted to mean condensation of intermediates on two separate proteins or built up via “waiting positions”^5^. Consistent with previous findings, tandem T domains in thalassospiramide assembly lines increase flux but are not mechanistic determinants of pass-back chain extension. This result is also consistent with the observation that the thalassospiramide assembly line from *Oceanibaculum pacificum* contains only a single T domain in module 4 and can still produce thalassospiramide A (**1**)^13^. There is no evidence that additional T domains promote “stalling” to allow for additional tailoring reactions, as having a single T domain in module 4 did not change product distribution but decreased levels of all analogs equally. We propose that thalassospiramide assembly lines adopt a conformational change to promote backward or forward passage of intermediates based on chain length. The assembly lines can also flexibly catalyze pass-back from module 4 to 2 and module 5 to 3.

Our work was facilitated by new methodology combining oligo recombineering with CRISPR-Cas9 counter selection for facile editing of large DNA constructs. Although our method was used solely for domain inactivation, it can be easily applied to perform other forms of multi-modular assembly line engineering, for example to change A domain specificity^33, 34^, to try to generate alternative chemical products.

It is curious that although the early stages of thalassospiramide biosynthesis are flexible, resulting in production of lipopeptides with a high degree of N-terminal structural diversity, the final stages are rather fixed. The C-terminal cyclic depsipeptide core of all thalassospiramide analogs is highly conserved, particularly the 12- membered ring and the α,β-unsaturated carbonyl moiety that together form the pharmacophore responsible for calpain protease inhibition^35^. Thus, the assembly line constructs a series of chemical products in which the structural elements that confer activity are maintained, while accessory elements such as the fatty acid and linear chain composition and length, which may confer target specificity, are variable. It has previously been speculated that biosynthetic promiscuity resulting in chemical diversity may be evolutionarily advantageous^36, 37^. Furthermore, the type of combinatorial biosynthesis observed in this study expands the portfolio of small molecules produced without introducing new assembly line modules or domains, or even additional tailoring enzymes, thus representing a means of expanding chemical diversity while minimizing genomic space. Currently, we cannot predict the plasticity of thalassospiramide assembly lines based on their amino acid sequence alone, and we imagine that biosynthetic multitasking within modular assembly lines may be more widespread, particularly if it does indeed confer evolutionary benefits.

Lastly, the flexibility of thalassospiramide assembly lines raises many questions regarding the overall structure of multi-modular enzymes. While structures of entire PKS^38–40^ and NRPS^41^ modules have been solved and structural features of multi-modular assembly lines have been proposed^24^, current models are difficult to reconcile with the flexibility that must accompany the various inter-module domain interactions we provide evidence for. Future efforts to understand the specific elements that enable and control assembly line flexibility will hopefully enhance efforts to engineer assembly line biosynthetic pathways.

## Acknowledgements

The authors thank J.R. van der Meer (University of Lausanne) for providing plasmid pRMR6K-Gm, V. de Lorenzo (National Center for Biotechnology-CSIC) for providing strain *P. putida* EM383, and D.L. Court (National Cancer Institute, NIH) for providing strain *E. coli* HME68. We are grateful to A. Edlund (J. Craig Venter Institute), P.Y. Qian (Hong Kong University of Science and Technology), H. Xia (Shanghai Institutes for Biological Sciences, CAS), and W. Fenical and P.R. Jensen (Scripps Institution of Oceanography, UCSD) for facilitating access to equipment, chemical standards, and bacterial strains. We also thank Y. Kudo, P.A. Jordan, J.R. Chekan, L.T. Hoang, and S. Carreto for helpful discussion and technical assistance. The research reported has been supported by National Institutes of Health grants F31-AI129299 to J.J.Z. and R01-GM085770 to B.S.M.

## Author contributions

J.J.Z. and B.S.M. designed the study. J.J.Z., X.T., T.H., and A.C.R. performed the experiments. J.J.Z. and B.S.M. analyzed the data and wrote the manuscript with input from all authors.

## Methods

### General methods

A complete list of the primers, plasmids, and strains used in this study can be found in **Supplementary Table 1**. DNA fragments larger than 3 kb were amplified with PrimeSTAR Max (Clontech Laboratories, Inc.); all other PCR products were amplified with PrimeSTAR HS DNA polymerase (Clontech Laboratories, Inc.). DNA isolations and manipulations were carried out using standard protocols. *Thalassospira* sp. CNJ-328 and *T. mobilis* KA081020-065 were grown in GYP media (glucose 10 g/L, yeast extract 4 g/L, peptone 2 g/L, sea salt 25 g/L). *S. cerevisiae* VL6-48N was grown in YPDA media (yeast extract 10 g/L, peptone 20 g/L, dextrose 20 g/L, adenine 100 mg/L) or selective histidine drop-out media containing 5-FOA (yeast nitrogen base without amino acids and ammonium sulfate 1.7 g/L, yeast synthetic dropout medium without histidine 1.9 g/L, sorbitol 182 g/L, dextrose 20 g/L, ammonium sulfate 5 g/L, adenine 100 mg/L, 5-FOA 1 g/L). *E. coli* and *P. putida* strains were grown in LB. *E. coli* TOP10 and DH5α λpir were used for standard cloning procedures. *E. coli* BW25113/pIJ790 was used for λ Red PCR targeting, and *E. coli* HME68 was used for oligo recombineering and CRISPR-Cas9 counter selection. *P. putida* EM383 was used for heterologous expression. All strains were grown at 30 °C except TOP10 and DH5α λpir, which were grown at 37 °C. Liquid cultures were grown shaking at 220 r.p.m. When necessary, *E. coli* cultures were supplemented with the following antibiotics: 50 µg/mL kanamycin (150 µg/mL for *P. putida*), 10 µg/mL gentamycin (30 µg/ml for *P. putida*), 50 µg/mL apramycin, 100 µg/mL ampicillin, 25 µg/mL chloramphenicol.

### Cloning and heterologous expression of *ttc* and *ttm*

The overall workflow for genetic manipulation and heterologous expression of the *ttc* and *ttm* pathways is outlined in **Supplementary Fig. 1**. Biosynthetic gene clusters were cloned from genomic DNA using a TAR cloning protocol described previously^42^. Cluster-specific capture vectors were generated through a one-step PCR amplification of pCAP-BAC (pCB) using primers pCB- ttcCV_F/R for *ttc* and pCB-ttmCV_F/R for *ttm*. Yeast clones were screened, and PCR positive constructs were purified and transferred to *E. coli* TOP10 for verification by restriction digestion. pCB was miniprepped from at least 25 mL of VL6-48N and at least 10 mL of TOP10. pJZ001, containing the *intB13* cassette, was cloned by Gibson Assembly using four PCR fragments (amplified using primers CEN6/ARS4_608F/R, intB13_2330F/R, aacC1_1257F/R, and pADH_597F/R) and the pACYCDuet-1 vector backbone linearized using HindIII and XhoI. Fully assembled pJZ001 was digested using HindIII and XhoI, and the 5155 bp fragment was gel purified and used for λ Red recombination to knock-in the *intB13* cassette into the pCB vector backbone, replacing several yeast genes no longer necessary. Retrofitted constructs were then transferred to *P. putida* by electroporation as described previously^15^ and selected for using kanamycin and gentamicin. Int-B13-mediated integration of *ttc* into the genome of *P. putida* was characterized by PCR and chemical analysis (**Supplementary Fig. 2**). Edited constructs were similarly introduced to *P. putida* and then tested for heterologous production of lipopeptide products.

### Extraction and LC-MS analysis

Precultures were inoculated with colonies picked from plates and grown overnight before being inoculated into full 50 mL cultures in 250 mL Erlenmeyer flasks (in triplicate). Full cultures were grown for 5 hours before addition of 1.5 g of autoclaved XAD7HP resin per 50 mL of culture. After 24 hours of additional incubation, culture ODs were measured and recorded at 600 nm and supernatant and cells were decanted. Resin was washed three times with Milli-Q water before extraction with 20 mL of ethyl acetate. Extracts were dried under nitrogen, resuspended in 200 µL of methanol, and filtered through a 0.22 µm filter prior to LC-MS-MS analysis.

An Agilent 1100 series HPLC system (Palo Alto, CA, U.S.A.) was coupled to a Bruker Impact II Q-TOF mass spectrometer ((Billerica, MA, U.S.A.) for LC-MS analysis. An Agilent ZORBAX 300SB-C18 LC column (300 Å, 5 µm, 150 × 0.5 mm) was used for LC separation. Mobile phase A was H_2_O in 0.1% FA and mobile phase B was ACN in 0.1% FA. The LC gradient was: t=0.00 min, 70%A; t=3.00 min, 70%A; t=5.00 min, 57%A; t=35.00 min, 57%A; t=49.00 min, 20%A; t=51.00 min, 0%A; t=52.00 min, 0%A; t=57.00 min, 70%A; t=60.00 min, 70%A. A post time of 10 min was set to re-equilibrate the column. For shorter runs, the LC gradient was: t=0.00 min, 70%A; t=3.00 min, 70%A; t=23.00 min, 20%A; t=24.00 min, 0%A; t=27.00 min, 70%A; t=30.00 min, 70%A. A post time of 3 min was set to re-equilibrate the column. Flow rate was 20 µL/min. Sample injection volume was 2 µL.

MS conditions for MS/MS spectra generation were set as follows: capillary voltage, 4500; nebulizer gas flow, 0.8 Bar; dry gas, 5.0 L/min at 180 °C; funnel 1 RF 150 Vpp; funnel 2 RF, 300 Vpp; isCID energy, 0 eV; hexapole RF: 50 Vpp; Quadrupole ion energy, 4 eV; low mass 50 m/z; collision cell energy, 20 – 50 eV; pre pulse storage 5.0 µs; collision RF, ramp from 350 to 800 Vpp; transfer time ramp from 50 to 100 µs; detection mass range 25 to 1000 m/z; MS2 spectra collection rate was 2.0 Hz.

All samples used for comparison were analyzed at the same time and under the same conditions. Values were normalized by culture ODs and compared only for peaks with identical MS spectra and retention time. Full MS spectra of thalassospiramide analogs not previously reported^11–13^ are shown in **Supplementary Fig. 10-23**.

### Gene deletion and complementation experiments

Gene deletions were made using λ Red PCR targeting as described previously^43^. Primers used to amplify the *aac(3)IV* cassette and confirm gene deletions are listed in **Supplementary Table 1**. Deletions using this cassette were made after addition of the *intB13* cassette, which contains a gentamycin resistance gene, as the apramycin resistance gene *aac(3)IV* confers resistance to gentamycin in *E. coli*. For complementation, *ttcD* and *ttcC* were amplified using primers Tn7-ttcD_F/R and Tn7-ttcC_F/R, respectively, and cloned into the mini-Tn7 vector pUC18R6K-mini-Tn7T-Gm^18^. Cloned constructs were introduced to *P. putida* by electroporation along with the helper plasmid pTNS1 and selected for using gentamycin. Complemented *P. putida* clones were then made electrocompetent and pCB constructs were transferred by electroporation and selected for using kanamycin and gentamycin. For complementation of TtcC- G234D (MT_6_ inactivation), the mutation was generated by amplification of pTn7::*ttcC* using primers Tn7-ttcC- g702a_F/R and confirmed by sequencing. Although a *ttcA* deletion construct was generated and a mini-Tn7 vector containing *ttcA* was prepared, the latter ultimately could not be transferred to *P. putida*, as no clones were obtained even after multiple attempts, likely due to the large size of the gene (>6.5 kb).

### Inactivation and testing of assembly line enzymatic domains

Assembly line domain active sites^19^ were identified using sequence alignments as illustrated in **Supplementary Fig. 8**. pJZ002, an ampicillin resistant version of the pCas9^20^ vector, was constructed as follows. First, the BsaI restriction site was first removed from the ampicillin resistance gene *bla* via PCR amplification of pKD20 using primers pKD20-g848a_F/R. The resulting construct was PCR amplified using primers ts-repA101_F/R and combined with a fragment amplified from pCas9 using primers pCas9_5058F/R by Gibson assembly. Spacer sequences were cloned into pJZ002 as described previously^20^. Spacer sequences and targeting oligos used to target specific domains are listed in **Supplementary Table 1**. Oligo recombination and CRISPR-Cas9 counter selection were performed as described previously^20, 44^, with several modifications. pCB-*ttc*-*int* or pCB-*ttm*-*int* was first transferred to *E. coli* HME68 by electroporation and selected for using kanamycin. A transformant was picked and grown at 30 °C to OD_600_ 0.4-0.5 and heat shocked for 15 minutes at 42 °C in a shaking water bath before being chilled on ice for 10 minutes. The cells were pelleted and washed with ice-cold water before being resuspended in a small volume of ice-cold water. 100 ng of pJZ002 containing the appropriate spacer was mixed with 100 ng of targeting oligo and the DNA mixture was then introduced to the cells prior to electroporation at 2.5 kV in a 2 mm gap electroporation cuvette. Cells were recovered for 2 hours at 30 °C shaking and plated on LB with kanamycin and ampicillin. Four clones of each mutant were picked, miniprepped, and screened by sequencing. If no correct mutant was identified, a new spacer sequence was designed and cloned into pJZ002 and the method was retried. Very subtle mutations could be recovered efficiently using an effective spacer sequence, although the effectiveness of the spacer could only be determined empirically. In total, four of 12 spacer sequences were redesigned to achieve successful editing (**Supplementary Table 1**). Correctly edited constructs were transferred to TOP10 and confirmed by restriction digestion (**Supplementary Fig. 9**) before being transferred to *P. putida* for heterologous expression experiments.

## References

1 Weissman, K. J. The structural biology of biosynthetic megaenzymes. Nat. Chem. Biol. 11, 660–670, doi:10.1038/nchembio.1883 (2015).

2 Cane, D. E. Programming of erythromycin biosynthesis by a modular polyketide synthase. J. Biol. Chem. 285, 27517–27523, doi:10.1074/jbc.R110.144618 (2010).

3 Robbel, L. & Marahiel, M. A. Daptomycin, a bacterial lipopeptide synthesized by a nonribosomal machinery. J. Biol. Chem. 285, 27501–27508, doi:10.1074/jbc.R110.128181 (2010).

4 Helfrich, E. J. & Piel, J. Biosynthesis of polyketides by trans-AT polyketide synthases. Nat. Prod. Rep. 33, 231–316, doi:10.1039/c5np00125k (2016).

5 Sussmuth, R. D. & Mainz, A. Nonribosomal peptide synthesis-principles and prospects. Angew. Chem. Int. Ed. Engl. 56, 3770–3821, doi:10.1002/anie.201609079 (2017).

6 Magarvey, N. A., Haltli, B., He, M., Greenstein, M. & Hucul, J. A. Biosynthetic pathway for mannopeptimycins, lipoglycopeptide antibiotics active against drug-resistant gram-positive pathogens. Antimicrob. Agents Chemother. 50, 2167–2177, doi:10.1128/AAC.01545-05 (2006).

7 Felnagle, E. A., Rondon, M. R., Berti, A. D., Crosby, H. A. & Thomas, M. G. Identification of the biosynthetic gene cluster and an additional gene for resistance to the antituberculosis drug capreomycin. Appl. Environ. Microbiol. 73, 4162–4170, doi:10.1128/AEM.00485-07 (2007).

8 Thomas, M. G., Chan, Y. A. & Ozanick, S. G. Deciphering tuberactinomycin biosynthesis: isolation, sequencing, and annotation of the viomycin biosynthetic gene cluster. Antimicrob. Agents Chemother. 47, 2823–2830 (2003).

9 Du, L., Sanchez, C., Chen, M., Edwards, D. J. & Shen, B. The biosynthetic gene cluster for the antitumor drug bleomycin from *Streptomyces verticillus* ATCC15003 supporting functional interactions between nonribosomal peptide synthetases and a polyketide synthase. Chem. Biol. 7, 623–642 (2000).

10 Gehring, A. M. et al. Iron acquisition in plague: modular logic in enzymatic biogenesis of yersiniabactin by *Yersinia pestis*. Chem. Biol. 5, 573–586 (1998).

11 Oh, D. C., Strangman, W. K., Kauffman, C. A., Jensen, P. R. & Fenical, W. Thalassospiramides A and B, immunosuppressive peptides from the marine bacterium *Thalassospira* sp. Org. Lett. 9, 1525–1528, doi:10.1021/ol070294u (2007).

12 Ross, A. C. et al. Biosynthetic multitasking facilitates thalassospiramide structural diversity in marine bacteria. J. Am. Chem. Soc. 135, 1155–1162, doi:10.1021/ja3119674 (2013).

13 Zhang, W. et al. Family-wide structural characterization and genomic comparisons decode the diversity-oriented biosynthesis of thalassospiramides by marine Proteobacteria. J. Biol. Chem 291, 27228–27238, doi:10.1074/jbc.M116.756858 (2016).

14 Xu, Y. et al. Bacterial biosynthesis and maturation of the didemnin anti-cancer agents. J. Am. Chem. Soc. 134, 8625–8632, doi:10.1021/ja301735a (2012).

15 Zhang, J. J., Tang, X., Zhang, M., Nguyen, D. & Moore, B. S. Broad-host-range expression reveals native and host regulatory elements that influence heterologous antibiotic production in Gram-negative bacteria. MBio 8, doi:10.1128/mBio.01291-17 (2017).

16 Miyazaki, R. & van der Meer, J. R. A new large-DNA-fragment delivery system based on integrase activity from an integrative and conjugative element. Appl. Environ. Microbiol. 79, 4440–4447, doi:10.1128/AEM.00711-13 (2013).

17 Martinez-Garcia, E., Nikel, P. I., Aparicio, T. & de Lorenzo, V. *Pseudomonas* 2.0: genetic upgrading of *P. putida* KT2440 as an enhanced host for heterologous gene expression. Microb. Cell Fact. 13, 159, doi:10.1186/s12934-014-0159-3 (2014).

18 Choi, K. H. & Schweizer, H. P. mini-Tn7 insertion in bacteria with single attTn7 sites: example *Pseudomonas aeruginosa*. Nat. Protoc. 1, 153–161, doi:10.1038/nprot.2006.24 (2006).

19 Marahiel, M. A., Stachelhaus, T. & Mootz, H. D. Modular peptide synthetases involved in nonribosomal peptide synthesis. Chem. Rev. 97, 2651–2674 (1997).

20 Jiang, W., Bikard, D., Cox, D., Zhang, F. & Marraffini, L. A. RNA-guided editing of bacterial genomes using CRISPR-Cas systems. Nat. Biotechnol. 31, 233–239, doi:10.1038/nbt.2508 (2013).

21 Rausch, C., Hoof, I., Weber, T., Wohlleben, W. & Huson, D. H. Phylogenetic analysis of condensation domains in NRPS sheds light on their functional evolution. BMC Evol. Biol. 7, 78, doi:10.1186/1471-2148-7-78 (2007).

22 Crosby, J. & Crump, M. P. The structural role of the carrier protein--active controller or passive carrier. Nat. Prod. Rep. 29, 1111–1137, doi:10.1039/c2np20062g (2012).

23 Calderone, C. T., Bumpus, S. B., Kelleher, N. L., Walsh, C. T. & Magarvey, N. A. A ketoreductase domain in the PksJ protein of the bacillaene assembly line carries out both alpha- and beta-ketone reduction during chain growth. Proc. Natl. Acad. Sci. U S A 105, 12809–12814, doi:10.1073/pnas.0806305105 (2008).

24 Marahiel, M. A. A structural model for multimodular NRPS assembly lines. Nat. Prod. Rep. 33, 136–140, doi:10.1039/c5np00082c (2016).

25 Bozhuyuk, K. A. J. et al. *De novo* design and engineering of non-ribosomal peptide synthetases. Nat. Chem. 10, 275–281, doi:10.1038/nchem.2890 (2018).

26 Awakawa, T. et al. Salinipyrone and pacificanone are biosynthetic by-products of the rosamicin polyketide synthase. Chembiochem 16, 1443–1447, doi:10.1002/cbic.201500177 (2015).

27 Moss, S. J., Martin, C. J. & Wilkinson, B. Loss of co-linearity by modular polyketide synthases: a mechanism for the evolution of chemical diversity. Nat. Prod. Rep. 21, 575–593, doi:10.1039/b315020h (2004).

28 Thomas, I., Martin, C. J., Wilkinson, C. J., Staunton, J. & Leadlay, P. F. Skipping in a hybrid polyketide synthase. Evidence for ACP-to-ACP chain transfer. Chem. Biol. 9, 781–787 (2002).

29 Wenzel, S. C., Meiser, P., Binz, T. M., Mahmud, T. & Muller, R. Nonribosomal peptide biosynthesis: point mutations and module skipping lead to chemical diversity. Angew. Chem. Int. Ed. Engl. 45, 2296–2301, doi:10.1002/anie.200503737 (2006).

30 Mootz, H. D. et al. Decreasing the ring size of a cyclic nonribosomal peptide antibiotic by in-frame module deletion in the biosynthetic genes. J. Am. Chem. Soc. 124, 10980–10981 (2002).

31 Gao, L. et al. Module and individual domain deletions of NRPS to produce plipastatin derivatives in *Bacillus subtilis*. Microb. Cell Fact. 17, 84, doi:10.1186/s12934-018-0929-4 (2018).

32 Yu, D., Xu, F., Zhang, S. & Zhan, J. Decoding and reprogramming fungal iterative nonribosomal peptide synthetases. Nat. Commun. 8, 15349, doi:10.1038/ncomms15349 (2017).

33 Davidsen, J. M. & Townsend, C. A. In vivo characterization of nonribosomal peptide synthetases NocA and NocB in the biosynthesis of nocardicin A. Chem. Biol. 19, 297–306, doi:10.1016/j.chembiol.2011.10.020 (2012).

34 Thirlway, J. et al. Introduction of a non-natural amino acid into a nonribosomal peptide antibiotic by modification of adenylation domain specificity. Angew. Chem. Int. Ed. Engl. 51, 7181–7184, doi:10.1002/anie.201202043 (2012).

35 Lu, L. et al. Mechanism of action of thalassospiramides, a new class of calpain inhibitors. Sci. Rep. 5, 8783, doi:10.1038/srep08783 (2015).

36 Fischbach, M. A. & Clardy, J. One pathway, many products. Nat. Chem. Biol. 3, 353–355, doi:10.1038/nchembio0707-353 (2007).

37 Firn, R. D. & Jones, C. G. Natural products--a simple model to explain chemical diversity. Nat. Prod. Rep. 20, 382–391 (2003).

38 Dutta, S. et al. Structure of a modular polyketide synthase. Nature 510, 512–517, doi:10.1038/nature13423 (2014).

39 Whicher, J. R. et al. Structural rearrangements of a polyketide synthase module during its catalytic cycle. Nature 510, 560–564, doi:10.1038/nature13409 (2014).

40 Edwards, A. L., Matsui, T., Weiss, T. M. & Khosla, C. Architectures of whole-module and bimodular proteins from the 6-deoxyerythronolide B synthase. J. Mol. Biol. 426, 2229–2245, doi:10.1016/j.jmb.2014.03.015 (2014).

41 Tanovic, A., Samel, S. A., Essen, L. O. & Marahiel, M. A. Crystal structure of the termination module of a nonribosomal peptide synthetase. Science 321, 659–663, doi:10.1126/science.1159850 (2008).

42 Zhang, J. J., Yamanaka, K., Tang, X. & Moore, B. S. Direct cloning and heterologous expression of natural product biosynthetic gene clusters by transformation-associated recombination. Methods Enzymol. (Accepted).

43 Tang, X. et al. Identification of thiotetronic acid antibiotic biosynthetic pathways by target-directed genome mining. ACS Chem. Biol. 10, 2841–2849, doi:10.1021/acschembio.5b00658 (2015).

44 Sawitzke, J. A. et al. Recombineering: highly efficient in vivo genetic engineering using single-strand oligos. Methods Enzymol. 533, 157–177, doi:10.1016/B978-0-12-420067-8.00010-6 (2013).

