## Supplementary material for "Avant-garde assembly-line biosynthesis expands diversity of cyclic lipodepsipeptide products": SI

**Table of contents**

**Supplementary Tables** p2-10

**Supplementary Figures** p11-33

**Supplementary Tables**

**Supplementary Table 1 l Primers, plasmids, and strains used in this study.**

a) Primers

| Name | Sequence |
|---|---|
| pCB-ttcCV_F | ATTTCCCCGAAAAGTGCCACCTGGGTCCTTTTCATCACGTGCTATAAAAAATGTCGAAAGCTACATATAAGGAACGT |
| pCB-ttcCV_R | TTTGCCGATTGGGCCGATTGATGCCGTGGCGGAGCCGCCGTACCAAACGCCCACTATTTATACCATGGGAGGC |
| pCB-ttmCV_F | CCATGTCCGCAAAAGTCGTGTTGACGATAGAGCTGGAACCGCAACTGCGTATGTCGAAAGCTACATATAAGGAACGT |
| pCB-ttmCV_R | ATCCGCGGCGTCGGTGCCGGCCGCGACGGTTCCGCCAAGGTCGCCGAGGTCCACTATTTATACCATGGGAGGC |
| CEN6/ARS4_608F | ATTCGAGCTCGGCGCGCCTGCAGGTCGACAAGCTTTCTATCGCCTTCTTGACGAG |
| CEN6/ARS4_608R | TGGCATCCATAGACCCACACCCAGCAGAATTCGGCCGCTTGCCTGTAACTTACAC |
| intB13_2330F | AGTTACAGGCAAGCGGCCGAATTCTGCTGGGTGTG |
| intB13_2330R | TTGTCACAACGCCGCTTGTGCAGGCGGCGCATATC |
| aacC1_1257F | CGCCGCCTGCACAAGCGGCGTTGTGACAATTTAC |
| aacC1_1257R | ACGGATACGATGGAGCGTTAGCCATGAGGGTTTAG |
| pADH_597F | CTCATGTTTAACGAACTAAACCCTCATGGCTAACGCTCCATCGTATCCGTATTCC |
| pADH_597R | AATTTCGCAGCAGCGGTTTCTTTACCAGACTCGAGTTTCGCGCAAGACGATTGAC |
| spacer-seq_F | CTGAAGTATATTTTAGATGA |
| d-ttc(-1)_F | **TTGGGACCGCATTCGCATCTTTCTGGCCGTCGCACGACAG**GGTCGGGCTGGGAAGTTCC |
| d-ttc(-1)_R | **GATCGGCGACGGCGGCGACCCCGCGGGTGTCCTTGGTATC**GTAGGCTGGAGCTGCTTCG |
| d-ttc(-1)-check_F | CGCAAAGCAGGTCGATTATG |
| d-ttc(-1)-check_R | CGTGCAGATACAGCAATCAG |
| d-ttc(+1)_F | **CATCGAAGAGACTGAAGAAGTGGCCGTCGCGAAGGCGGAT**GGTCGGGCTGGGAAGTTCC |
| d-ttc(+1)_R | **TCGCACCTGCTGCCGGTTTGGCAGTGGTTTTGCCGCCTTT**GTAGGCTGGAGCTGCTTCG |
| d-ttc(+1)-check_F | GATGATCAATCAGGCCTTGG |
| d-ttc(+1)-check_R | GTGTTGCACGTTGCTACTTC |
| d-ttc(+2) _F | **CCTGCTGGTCGCTATTTTCGCGTTGGCGCCGCAGATTTTC**GGTCGGGCTGGGAAGTTCC |
| d-ttc(+2) _R | **CGCCCCCGCTACCACTGGATTGATCAAAGACCGGCGCAAT**GTAGGCTGGAGCTGCTTCG |
| d-ttc(+2)-check_F | GCCACTTCTTCAGTCTCTTC |
| d-ttc(+2)-check_R | ATACACCCGACGACGAGATG |

| | |
|---|---|
| d-ttc(+3+4)_F | **GGGGATATTTGCCAATCTTGCCAAGGTGATGCCGGTTGTC**GGTCGGGCTGGGAAGTTCC |
| d-ttc(+3+4)_R | **AATCAAGGTCTTTAAGCTTTTCACGAAGCTCCCGCGACGG**GTAGGCTGGAGCTGCTTCG |
| d-ttc(+3+4)-check_F | ATGCTGATCGCCCGTCAGTC |
| d-ttc(+3+4)-check_R | TCCGTGATCGCGGCCTTAAC |
| d-ttcD_F | **CCCGAAGGGCACCGGCACATTTGTGTCGTCCGATATCCTG**GGTCGGGCTGGGAAGTTCC |
| d-ttcD_R | **GATCCGGCAATGTCAGGCCAAGCCACCCCTGCGCACCGGG**GTAGGCTGGAGCTGCTTCG |
| d-ttcD-check_F | CGAAGGCGATCAATATAGCG |
| d-ttcD-check_R | GCGTCCGAATTTGTCAAAGG |
| Tn7-ttcD_F | TTTGAAGCTAATTCGATCATGCATGAGCTCATAGCGGCTGGCAAAGGGTC |
| Tn7-ttcD_R | GGTTGGCCTGCAAGGCCTTCGCGAGGTACCGCAAGGCGTCCGAATTTGTC |
| d-ttcC_F | **CATCGCAACCCGCGATGAAACCGGCGAAATCAGCTATACT**GGTCGGGCTGGGAAGTTCC |
| d-ttcC_R | **TCCGGCAAACCTGCCCTGGGCCTGATTTTTCAGTCCATCA**GTAGGCTGGAGCTGCTTCG |
| d-ttcC-check_F | GAATGGCACGCCCGATGTTC |
| d-ttcC-check_R | GGTTCGATCCGGCAAACCTG |
| Tn7-ttcC_F | TTTGAAGCTAATTCGATCATGCATGAGCTCTCTGTTCCTGACCGAAGAAG |
| Tn7-ttcC_R | GGTTGGCCTGCAAGGCCTTCGCGAGGTACCTGCCAATTTCGGGCAGTATC |
| Tn7-ttcC-g702a_F | TGCCCGGGTGCTTGAAATCG**A**CTGCGCGTCCGGCTTCACCA |
| Tn7-ttcC-g702a_R | TGGTGAAGCCGGACGCGCAG**T**CGATTTCAAGCACCCGGGCA |
| ttcC-g702a-check_F | AGTGCCTTTACCGGGCAACC |
| ttcC-g702a-check_R | CTGCCGACCGATGACATGTG |
| d-ttcA_F | **ATTTCAAAAACCGCTTGCTGCATGGTGCAATCAACGTTTG**GGTCGGGCTGGGAAGTTCC |
| d-ttcA_R | **TGGTAGGCGCGCTGTTTTTATCAAAGGATGCAATCAGGTT**GTAGGCTGGAGCTGCTTCG |
| d-ttcA-check_F | AGGCACCTAAACAGCGTACC |
| d-ttcA-check_R | TCCCTGTTCGCGTGTCGAAG |
| Tn7-ttcA_F | TTTGAAGCTAATTCGATCATGCATGAGCTCCTGTTCAGGCACCTAAACAG |
| Tn7-ttcA_R | GGTTGGCCTGCAAGGCCTTCGCGAGGTACCCATGCAGGCTTAGCACAATG |
| pKD20-g848a_F | TCTGGAGCCGGTGAGCGTGG**A**TCTCGCGGTATCATTGCAGC |
| pKD20-g848a_R | GCTGCAATGATACCGCGAGA**T**CCACGCTCACCGGCTCCAGA |
| bla*-check_214F | GATGGAGGCGGATAAAGTTG |
| bla*-check_214R | ACCTATCTCAGCGATCTGTC |
| ts-repA101_F | **CAAAAGAAGAGTAGTGTGATCGTCCATTCC**TAAGCTAGCCCATGGGTATGGA |
| ts-repA101_R | **TATCATTCTACATTTAGGCGCTGCCATCTT**CAGGTGGCACTTTTCGGGGA |

| | |
|---|---|
| pCas9_5058F | AAGATGGCAGCGCCTAAATG |
| pCas9_5058R | GGAATGGACGATCACACTAC |
| ttcA-C1a_H154A_F | CAAGGGACGTTTCTGGTGGGTCCGGGTGTATCA**CGCA**CT**C**GTCTGTGACGGATATGCAGGCCATCTGATG |
| ttcA-C1a-spacer_F | AAACGTTTCTGGTGGGTCCGGGTGTATCATCATCG |
| ttcA-C1a-spacer_R | AAAACGATGATGATACACCCGGACCCACCAGAAAC |
| ttcA-C1a-check_232F | GGTTCTGATCGCCCATAACG |
| ttcA-C1a-check_232R | GAATGTGCTTTCGGGCACTG |
| ttcA-A1a_G631D_F | GTGACGATATCGCCTTTGTCTTCCATACGTCGG**ACAGC**ACCGGGCAACCAAAACCGGTTCCCGTTCATCA |
| ttcA-A1a(G)-spacer_F | AAACTGACGATATCGCCTTTGTCTTCCATACGTCG |
| ttcA-A1a(G)-spacer_R | AAAACGACGTATGGAAGACAAAGGCGATATCGTCA |
| ttcA-A1a(G)-check_380F | GATTGCCGACTGGTTCTGAG |
| ttcA-A1a(G)-check_380R | TGGGTCATCAGCCACATACG |
| ttcA-A1a_K972A_F | TGGGTGGAAACGCTTCCTTTGCTACCGTC**C**GG**GGCG**AT**A**GA**C**CGCAAGGCACTTGCCCAATTTGCTCAGG |
| ttcA-A1a(K)-spacer_F | AAACGGCAAGTGCCTTGCGATCGATTTTGCCAGAG |
| ttcA-A1a(K)-spacer_R | AAAACTCTGGCAAAATCGATCGCAAGGCACTTGCC |
| ttcA-A1a(K)-check_235F | GACGGAAGCCATTCGAAACG |
| ttcA-A1a(K)-check_235R | TGCGCATCTGATCGGGTTTG |
| ttcA-T1a_S1041A_F | TCGATACCAACCTGTTTGAAGCTGGCGCCCA**CG**C**CC**T**C**CTGGTACCGCGTGCACAGTTTGCCCTGTCAAA |
| ttcA-T1a-spacer_F | AAACGCAAACTGTGCACGCGGTACCAGCAATGAAG |
| ttcA-T1a-spacer_R | AAAACTTCATTGCTGGTACCGCGTGCACAGTTTGC |
| ttcA-T1a-check_297F | CAAACAGCCTGTTGCACAAG |
| ttcA-T1a-check_297R | TCGGGAACTGTCTGGTTATC |
| ttcA-KS1b_A1273G C1274A_F | CCTGACCGGCCCAGCCGTCGCGTCCTCGACCG**GTGCG**TCGACCGGCCTTGTCAATATTGCGCTTGCCGTC |
| ttcA-KS1b-spacer_F | AAACCAAGCGCAATATTGACAAGGCCGGTCGAACG |
| ttcA-KS1b-spacer_R | AAAACGTTCGACCGGCCTTGTCAATATTGCGCTTG |
| ttcA-KS1b-check_293F | TTGGTTTCCCGACCTATCTG |
| ttcA-KS1b-check_293R | GCCCTGCGTGAAGTAATATC |
| ttcA-AT1b_S1728A_F | CGGCATTTCACCCGCCGCGCTGGCCGGGCAC**GCA**ATTGGCGAATATGTCGCAGCCTGCATTGGCGGGGTC |
| ttcA-AT1b-spacer_F | AAACTTCACCCGCCGCGCTGGCCGGGCACAGCATG |
| ttcA-AT1b-spacer_R | AAAACATGCTGTGCCCGGCCAGCGCGGCGGGTGAA |
| ttcA-AT1b-spacer2_F | AAACAGGCTGCGACATATTCGCCAATGCTGTGCCG |

| | |
|---|---|
| ttcA-AT1b-spacer2_R | AAAACGGCACAGCATTGGCGAATATGTCGCAGCCT |
| ttcA-AT1b-check_283F | AACATCAAGCCGCATGGCAC |
| ttcA-AT1b-check_283R | AAAGGCCGACAGCAGATCCC |
| ttcB-A3_K2045A_F | CCCTTGATCATCTCCCGCTTACCAGC**TCA**GG**TGCA**GT**G**GACCGAAAGGCCCTTTCGGGCACACCGATGGC |
| ttcB-A3-spacer_F | AAACTTGATCATCTCCCGCTTACCAGCAGCGGCAG |
| ttcB-A3-spacer_R | AAAACTGCCGCTGCTGGTAAGCGGGAGATGATCAA |
| ttcB-A3-spacer2_F | AAACAAAGGGCCTTTCGGTCGACCTTGCCGCTGCG |
| ttcB-A3-spacer2_R | AAAACGCAGCGGCAAGGTCGACCGAAAGGCCCTTT |
| ttcB-A3-check_208F | AGGCGACCTGACCATTGGTG |
| ttcB-A3-check_208R | ATCGGGCTTTGCCGCGATAG |
| ttcB-T4a_S3509V_F | CTCGGATGATGATTTCTTTGCTCTTGGCGG**T**GA**CGTT**AT**T**AC**A**GGCATGCAGATCGTCGACCGCATCAAT |
| ttcB-T4a-spacer_F | AAACTGATGCGGTCGACGATCTGCATGCCGGTTAG |
| ttcB-T4a-spacer_R | AAAACTAACCGGCATGCAGATCGTCGACCGCATCA |
| ttcB-T4b_S3595V_F | CGGACGAGGATTTCTATGCTTTGGGCGG**T**GA**CGTT**AT**T**AC**A**GGCATGCAGATCGTTGATCGCATGAATGC |
| ttcB-T4b-spacer_F | AAACTCATGCGATCAACGATCTGCATGCCGGTGAG |
| ttcB-T4b-spacer_R | AAAACTCACCGGCATGCAGATCGTTGATCGCATGA |
| ttcB-T4-check_566F | GTTTCCGCACCGGTATCTCC |
| ttcB-T4-check_566R | GGCGATGGTGTCGCTGTTTC |
| ttmA-C1_H163A_F | GCCCGATCGCCATCGCTGGATCCGCTGCTATCA**CGCC**CT**C**ATCCTGGATGGTCAGGGTGGCATGATCCTG |
| ttmA-C1-spacer_F | AAACGCTGGATCCGCTGCTATCATCATCTGATCCG |
| ttmA-C1-spacer_R | AAAACGGATCAGATGATGATAGCAGCGGATCCAGC |
| ttmA-C1-spacer2_F | AAACTGACCATCCAGGATCAGATGATGATAGCAGG |
| ttmA-C1-spacer2_R | AAAACCTGCTATCATCATCTGATCCTGGATGGTCA |
| ttmA-C1-check-194F | TTGCCCAAGAGGCCTTCGAG |
| ttmA-C1-check-194R | GATGAGCGCGGTGTAGATCC |
| ttmA-C2_H700A_F | GGACCAACCCGAGCGGTTGCGGGTGGTGGTCGAT**GCC**CTGGTCTTCGACGGCGAAAGCCGCACGGTGTTC |
| ttmA-C2-spacer_F | AAACCCGAGCGGTTGCGGGTGGTGGTCGATCATCG |
| ttmA-C2-spacer_R | AAAACGATGATCGACCACCACCCGCAACCGCTCGG |
| ttmA-C2-spacer2_F | AAACGCGGGTGGTGGTCGATCATCTGGTCTTCGAG |
| ttmA-C2-spacer2_R | AAAACTCGAAGACCAGATGATCGACCACCACCCGC |
| ttmA-C2-check_435F | ATGACGATGCGGCGGAAATG |

| | |
|---|---|
| ttmA-C2-check_435R | ACAGGGCGGTGGTTGCAATG |
| ttmB-MT6_G230D_F | TCACGCCCGCCAGCCGGGTGCTGGAAATCG**AT**TGCGCCTCGGGCTTCACGCTCCGTGCCC TGGCGCCGCT |
| ttmB-MT6-spacer_F | AAACCCAGCCGGGTGCTGGAAATCGGCTGCGCCTG |
| ttmB-MT6-spacer_R | AAAACAGGCGCAGCCGATTTCCAGCACCCGGCTGG |
| ttmB-MT6-check_555F | ATATCGCCCATCTCGTCTGC |
| ttmB-MT6-check_555R | CGCGGATATTGCCCAGAAAG |

b) Plasmids

| **Name** | **Description** | **Reference** |
|---|---|---|
| pCAP-BAC (pCB) | Yeast-E. coli artificial chromosome shuttle vector; HIS3 and Kan$^R$ | This study |
| pACR11 | pCAP01 carrying *ttc* | This study |
| pRMR6K-Gm | R6K replicon, Pcirc-lacO4-*intB13*, *oriT*, FRT, MCS, Gm$^R$ | 1 |
| pJZ001 | pACYCDuet-1 carrying *intB13* cassette for pCB knock-in | This study |
| pCas9 | *cas9* and *tracr* vector; Chl$^R$ | 2 |
| pKD20 | *repA101*(ts)-*araBp-gam-bet-exo-ori101-bla*; Amp$^R$ | 3 |
| pJZ002 | *repA101(ts)-cas9-tracr-bla** (lacking BsaI site)*;* Amp$^R$ | This study |
| pCAP05-*ttc* | pCAP05 carrying *ttc* (*-1* through *+6*) | This study |
| pCB-*ttc* | pCB carrying *ttc* | This study |
| pCB-*ttm* | pCB carrying *ttm* (*-1* through *+1*) | This study |
| pCB-*ttc-int* | pCB-*ttc* with *intB13* | This study |
| pCB-*ttm-int* | pCB-*ttm* with *intB13* | This study |
| pCB-*ttc-int*-Δ(*-1*) | pCB-*ttc-int* Δ(*ttc -1*) | This study |
| pCB-*ttc-int*-Δ(*+1*) | pCB-*ttc-int* Δ(*ttc +1*) | This study |
| pCB-*ttc-int*-Δ(*+2*) | pCB-*ttc-int* Δ(*ttc +2*) | This study |
| pCB-*ttc-int*-Δ(*+3+4*) | pCB-*ttc-int* Δ(*ttc +3*)(*ttc +4*) | This study |
| pCB-*ttc-int*-Δ(*ttcD*) | pCB-*ttc-int* Δ(*ttcD*) | This study |
| pTNS1 | R6K replicon; encodes the *tnsABC+D* specific transposition pathway; Amp$^R$ | 4 |
| pUC18R6K-mini-Tn7T-Gm | R6K replicon; mini-Tn7 vector; Amp$^R$, Gm$^R$ | 4 |
| pTn7::*ttcD* | pUC18R6K-mini-Tn7T-Gm carrying *ttcD* | This study |
| pCB-*ttc-int*-Δ(*ttcC*) | pCB-*ttc-int* Δ(*ttcC*) | This study |
| pTn7::*ttcC* | pUC18R6K-mini-Tn7T-Gm carrying *ttcC* | This study |
| pTn7::*ttcC*-g702a | pUC18R6K-mini-Tn7T-Gm carrying *ttcC*-g702a (G234D) | This study |
| pJZ002::*vioX* | pJZ002 with *vioX* spacer | This study |
| pJZ002::C1a | pJZ002 with C1a spacer | This study |
| pCB-*ttc-int*-C1a 1 | pCB-*ttc-int* with TtcA-H154A clone 1 | This study |
| pCB-*ttc-int*-C1a 2 | pCB-*ttc-int* with TtcA-H154A clone 2 | This study |
| pJZ002::A1a(G) | pJZ002 with A1a(G) spacer | This study |
| pCB-*ttc-int*-A1a (G631D) | pCB-*ttc-int* with TtcA-G631D | This study |
| pJZ002:A1a(K) | pJZ002 with A1a(K) spacer | This study |
| pCB-*ttc-int*-A1a (K972A) 1 | pCB-*ttc-int* with TtcA-K972A clone 1 | This study |
| pCB-*ttc-int*-A1a (K972A) 2 | pCB-*ttc-int* with TtcA-K972A clone 2 | This study |
| pJZ002::T1a | pJZ002 with T1a spacer | This study |
| pCB-*ttc-int*-T1a | pCB-*ttc-int* with TtcA-S1041A | This study |
| pJZ002::KS1b | pJZ002 with KS1b spacer | This study |
| pCB-*ttc-int*-KS1b | pCB-*ttc-int* with TtcA-A1273G-C1274A | This study |
| pJZ002:AT1b | pJZ002 with AT1b spacer 1 | This study |
| pJZ002:AT1b-2 | pJZ002 with AT1b spacer 2 | This study |
| pCB-*ttc-int*-AT1b | pCB-*ttc-int* with TtcA-S1728A | This study |

| | | |
|---|---|---|
| pJZ002:A3 | pJZ002 with A3 spacer 1 | This study |
| pJZ002:A3-2 | pJZ002 with A3 spacer 1 | This study |
| pCB-*ttc-int*-A3 | pCB-*ttc-int* with TtcB-K2045A | This study |
| pJZ002:T4a | pJZ002 with T4a spacer | This study |
| pCB-*ttc-int*-T4 A | pCB-*ttc-int* with TtcB-Δ3511-3596 version A | This study |
| pJZ002:T4b | pJZ002 with T4b spacer | This study |
| pCB-*ttc-int*-T4 B | pCB-*ttc-int* with TtcB-Δ3511-3596 version B | This study |
| pJZ002::C1 | pJZ002 with C1 spacer 1 | This study |
| pJZ002::C1-2 | pJZ002 with C1 spacer 2 | This study |
| pCB-*ttm-int*-C1 1 | pCB-*ttm-int* with TtmA-H163A clone 1 | This study |
| pCB-*ttm-int*-C1 2 | pCB-*ttm-int* with TtmA-H163A clone 2 | This study |
| pJZ002::C2 | pJZ002 with C2 spacer 1 | This study |
| pJZ002::C2-2 | pJZ002 with C2 spacer 2 | This study |
| pCB-*ttm-int*-C2 1 | pCB-*ttm-int* with TtmA-H700A clone 1 | This study |
| pCB-*ttm-int*-C2 2 | pCB-*ttm-int* with TtmA-H700A clone 2 | This study |
| pJZ002::MT6 | pJZ002 with MT6 spacer | This study |
| pCB-*ttm-int*-MT6 | pCB-*ttm-int* with TtmB-G230D | This study |

c) Strains

| **Name** | **Description** | **Reference** |
|---|---|---|
| *S. cerevisiae* VL6-48N | *MATα trp1-Δ1 ura3-Δ1 ade2-101 his3-Δ200 lys2 met14 cir°* | 5 |
| *E. coli* TOP10 | *mcrA, Δ(mrr-hsdRMS-mcrBC), Phi80lacZ(del)M15, ΔlacX74, deoR, recA1, araD139, Δ(ara-leu)7697, galU, galK, rpsL(SmR), endA1, nupG* | N/A |
| *E. coli* DH5α λpir | *sup E44, ΔlacU169 (ΦlacZΔM15), recA1, endA1, hsdR17, thi-1, gyrA96, relA1, λpir* | N/A |
| *E. coli* BW25113/ pIJ790 | Strain: Δ(araD-araB)567, ΔlacZ4787(::rrnB-4), lacIp-4000(lacIQ), λ-, rpoS369(Am), rph-1, Δ(rhaD-rhaB)568, hsdR514<br>Plasmid: [*oriR101*], [*repA101*(ts)], *araBp-gam-be-exo* | 6 |
| *E. coli* HME68 | W3110 *galKtyr145UAG ΔlacU169 [λ cI857 Δ(cro-bioA)] mutS<>cat* | 7 |
| *P. putida* EM383 | KT2440 derivative; Δprophage1 Δprophage4 Δprophage3 Δprophage2 Δ*Tn7* Δ*endA-1* Δ*endA-2* Δ*hsdRMS* Δflagellum Δ*Tn4652* Δ*recA* | 8 |

References:


1 Miyazaki, R. & van der Meer, J. R. A new large-DNA-fragment delivery system based on integrase activity from an integrative and conjugative element. *Appl. Environ. Microbiol.* **79,** 4440-4447, doi:10.1128/AEM.00711-13 (2013).

2 Jiang, W., Bikard, D., Cox, D., Zhang, F. & Marraffini, L. A. RNA-guided editing of bacterial genomes using CRISPR-Cas systems. *Nat. Biotechnol.* **31,** 233-239, doi:10.1038/nbt.2508 (2013).

3 Datsenko, K. A. & Wanner, B. L. One-step inactivation of chromosomal genes in Escherichia coli K-12 using PCR products. *Proc. Natl. Acad. Sci. U S A* **97,** 6640-6645, doi:10.1073/pnas.120163297 (2000).

4 Choi, K. H. & Schweizer, H. P. mini-Tn7 insertion in bacteria with single attTn7 sites: example *Pseudomonas aeruginosa*. *Nat. Protoc.* **1,** 153-161, doi:10.1038/nprot.2006.24 (2006).

5 Noskov, V. N. *et al.* A general cloning system to selectively isolate any eukaryotic or prokaryotic genomic region in yeast. *BMC Genomics* **4,** 16, doi:10.1186/1471-2164-4-16 (2003).

6 Gust, B., Challis, G. L., Fowler, K., Kieser, T. & Chater, K. F. PCR-targeted Streptomyces gene replacement identifies a protein domain needed for biosynthesis of the sesquiterpene soil odor geosmin. *Proc. Natl. Acad. Sci. U S A* **100,** 1541-1546, doi:10.1073/pnas.0337542100 (2003).

7 Sawitzke, J. A. *et al.* Recombineering: in vivo genetic engineering in *E. coli*, *S. enterica*, and beyond. *Methods Enzymol.* **421,** 171-199, doi:10.1016/S0076-6879(06)21015-2 (2007).

8 Martinez-Garcia, E., Nikel, P. I., Aparicio, T. & de Lorenzo, V. *Pseudomonas* 2.0: genetic upgrading of *P. putida* KT2440 as an enhanced host for heterologous gene expression. *Microb. Cell Fact.* **13,** 159, doi:10.1186/s12934-014-0159-3 (2014).

**Supplementary Table 2 | Annotation and BLAST results of biosynthetic pathway genes.**

*a) ttc*

| Gene | Size (aa) | Proposed function | Description (top BLAST hit) | Identity | NCBI accession |
|---|---|---|---|---|---|
| *ttc* -7 | 453 | Regulation | Sigma-54-dependent Fis family transcriptional regulator [Thalassospira] | 99% | WP_082824890.1 |
| *ttc* -6 | 149 | Hypothetical | TonB-dependent receptor [Thalassospira lucentensis] | 99% | WP_062953563.1 |
| *ttc* -5 | 478 | Transport | TonB-dependent receptor [Thalassospira lucentensis] | 99% | WP_062953563.1 |
| *ttc* -4 | 270 | Hypothetical | ferric iron reductase protein FhuF [Thalassospira xiamenensis] | 99% | SOC29973.1 |
| *ttc* -3 | 537 | Transport | ABC transporter ATP-binding protein/permease [Thalassospira xiamenensis] | 99% | WP_114109924.1 |
| *ttc* -2 | 499 | Other | methylmalonate-semialdehyde dehydrogenase (CoA acylating) [Thalassospira xiamenensis] | 99% | WP_062959055.1 |
| *ttc* -1 | 295 | Regulation | LysR family transcriptional regulator [Thalassospira xiamenensis] | 99% | WP_114109923.1 |
| *ttcA* | 2126 | NRPS/PKS | hybrid non-ribosomal peptide synthetase/type I polyketide synthase [Thalassospira xiamenensis] | 97% | WP_097053431.1 |
| *ttcB* | 5186 | NRPS/PKS | non-ribosomal peptide synthetase [Thalassospira lucentensis] | 96% | WP_062953557.1 |
| *ttcC* | 1174 | NRPS/PKS | alpha/beta fold hydrolase [Thalassospira xiamenensis] | 98% | WP_062959059.1 |
| *ttcD* | 253 | NRPS/PKS PPTase | hypothetical protein [Thalassospira lucentensis] | 95% | WP_062953555.1 |
| *ttc* +1 | 186 | Hypothetical | DUF697 domain-containing protein [Thalassospira sp. MCCC 1A03138] | 100% | WP_085646013.1 |
| *ttc* +2 | 420 | Hypothetical | Prohibitin family protein [Thalassospira] | 99% | WP_062953553.1 |

| | | | | | |
|---|---|---|---|---|---|
| *ttc* +3 | 63 | Hypothetical | hypothetical protein [Thalassospira] | 100% | WP_062953552.1 |
| *ttc* +4 | 79 | Hypothetical | DUF697 domain-containing protein [Thalassospira] | 100% | WP_085646011.1 |
| *ttc* +5 | 260 | Hypothetical | hypothetical protein [Thalassospira xiamenensis] | 99% | WP_097053437.1 |
| *ttc* +6 | 324 | Hypothetical | MFS transporter [Thalassospira xiamenensis] | 100% | WP_097053438.1 |

b) *ttm* (only *ttm-1* through *ttm +1* was cloned)

| Gene | Size (aa) | Proposed function | Description (top BLAST hit) | Identity | NCBI accession |
|---|---|---|---|---|---|
| *ttm* -6 | 1220 | Hypothetical | hypothetical protein [Erythrobacter sp. YT30] | 48% | WP_067603228.1 |
| *ttm* -5 | 423 | Other | type II toxin-antitoxin system HipA family toxin [Roseomonas rosea] | 86% | WP_073135195.1 |
| *ttm* -4 | 81 | Regulation | transcriptional regulator [Sandarakinorhabdus cyanobacteriorum] | 90% | WP_094472265.1 |
| *ttm* -3 | 707 | Other | catalase [Catalinimonas alkaloidigena] | 71% | WP_089678827.1 |
| *ttm* -2 | 163 | Other | CBS domain-containing protein [Mesorhizobium delmotii] | 66% | SJM34906.1 |
| *ttm* -1 | 242 | Other | uracil-DNA glycosylase [Sphingomonas sp. PR090111-T3T-6A] | 51% | WP_019834353.1 |
| *ttmA* | 5215 | NRPS/PKS | TtbA [Tistrella bauzanensis] | 63% | AGC65516.1 |
| *ttmB* | 1166 | NRPS/PKS | TtbB [Tistrella bauzanensis] | 69% | AGC65517.1 |
| *ttm* +1 | 189 | Other | HNH endonuclease [Defluviimonas alba] | 67% | WP_084739999.1 |
| *ttm* +2 | 104 | Other | antitoxin of toxin-antitoxin stability system [Rhizobium sp. 60-20] | 71% | OJY78535.1 |
| *ttm* +3 | 82 | Other | type II toxin-antitoxin system ParD family antitoxin [Neorhizobium galegae] | 75% | WP_038540781.1 |

| | | | | | |
|---|---|---|---|---|---|
| *ttm* +4 | 98 | Other | type II toxin-antitoxin system RelE/ParE family toxin [Phyllobacteriaceae bacterium SYSU D60010] | 48% | WP_119273623.1 |
| *ttm* +5 | 594 | Hypothetical | hypothetical protein AMS22_02705 [Thiotrichales bacterium SG8_50] | 37% | KPK56010.1 |
| *ttm* +6 | 304 | Regulation | LysR family transcriptional regulator [Paraburkholderia nodosa] | 69% | WP_051481188.1 |
| *ttm* +7 | 324 | Other | nitronate monooxygenase [Alcaligenes faecalis] | 80% | WP_083053698.1 |
| *ttm* +8 | 189 | Other | cytochrome b561 [Alcaligenes faecalis] | 71% | WP_026482775.1 |
| *ttm* +9 | 395 | Hypothetical | hypothetical protein A3D94_00820 [Alphaproteobacteria bacterium RIFCSPHIGHO2_12_FULL_66_14] | 37% | OFX06623.1 |

### Supplementary Figures

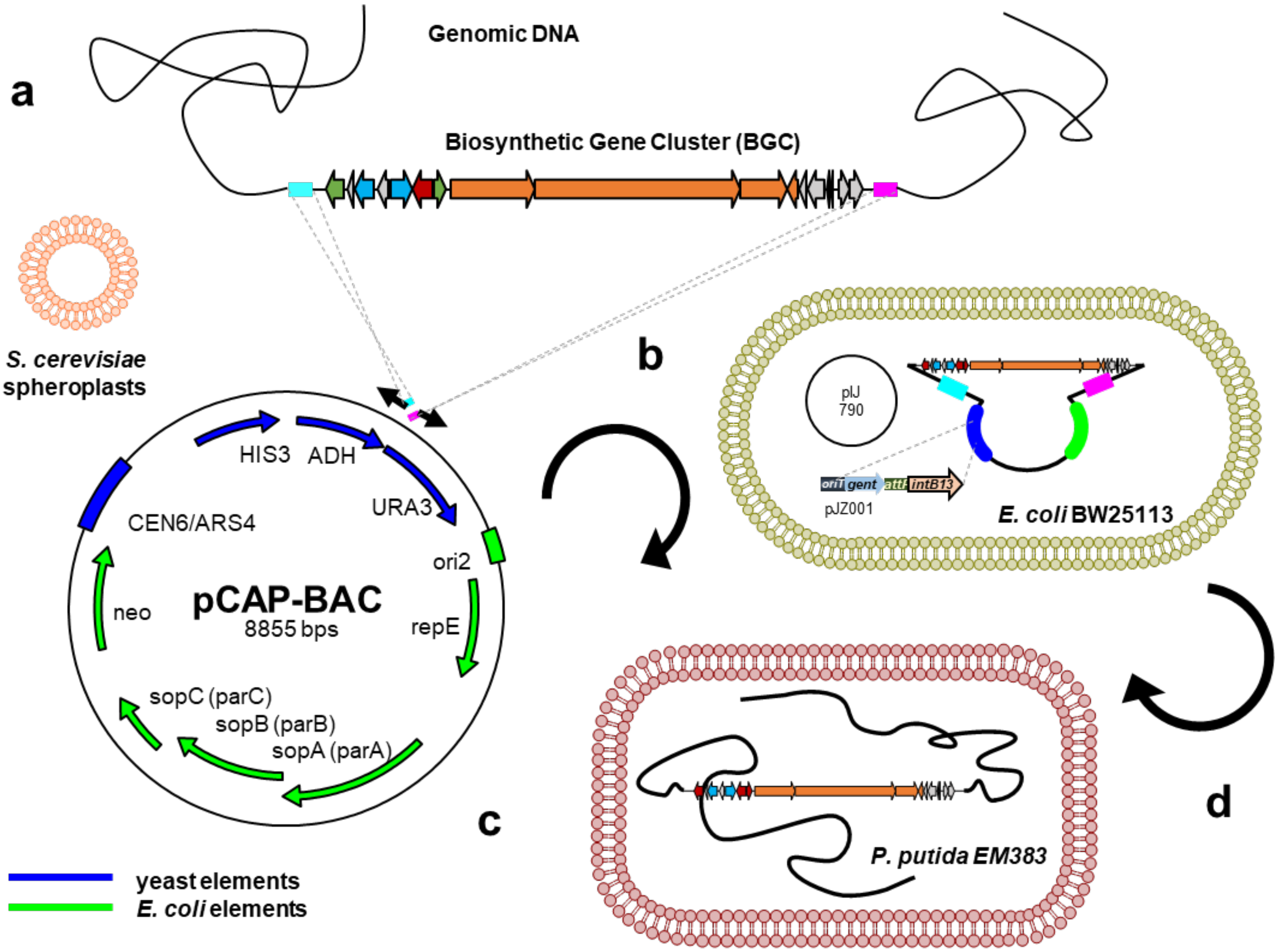


**Supplementary Fig. 1 | Platform for heterologous expression of thalassospiramide gene clusters**. **a**) Direct cloning of gene clusters from genomic DNA was facilitated by transformation-associated recombination (TAR) cloning in *Saccharomyces cerevisiae* using new vector pCAP-BAC (pCB), which can be assembled into a cluster-specific capture vector by a one-step PCR reaction using long primers with homology arms flanking the genomic loci of interest. **b**) Individual clones that have been picked and screened are first transferred to *E. coli* TOP10 for maintenance and confirmation. Then, recombineering strain *E. coli* BW25113/pIJ790 is used to introduce the *intB13* integration cassette (from pJZ001) into the pCB vector backbone via Lambda Red recombination. **c**) Constructs retrofitted with IntB13 are transferred and integrated into the genome of the heterologous host *P. putida* EM383, which is a derivate of KT2440. **d**) Cloned constructs can be readily manipulated using *E. coli* recombineering tools and re-introduced to the heterologous host to rapidly connect genotype to chemical phenotype.

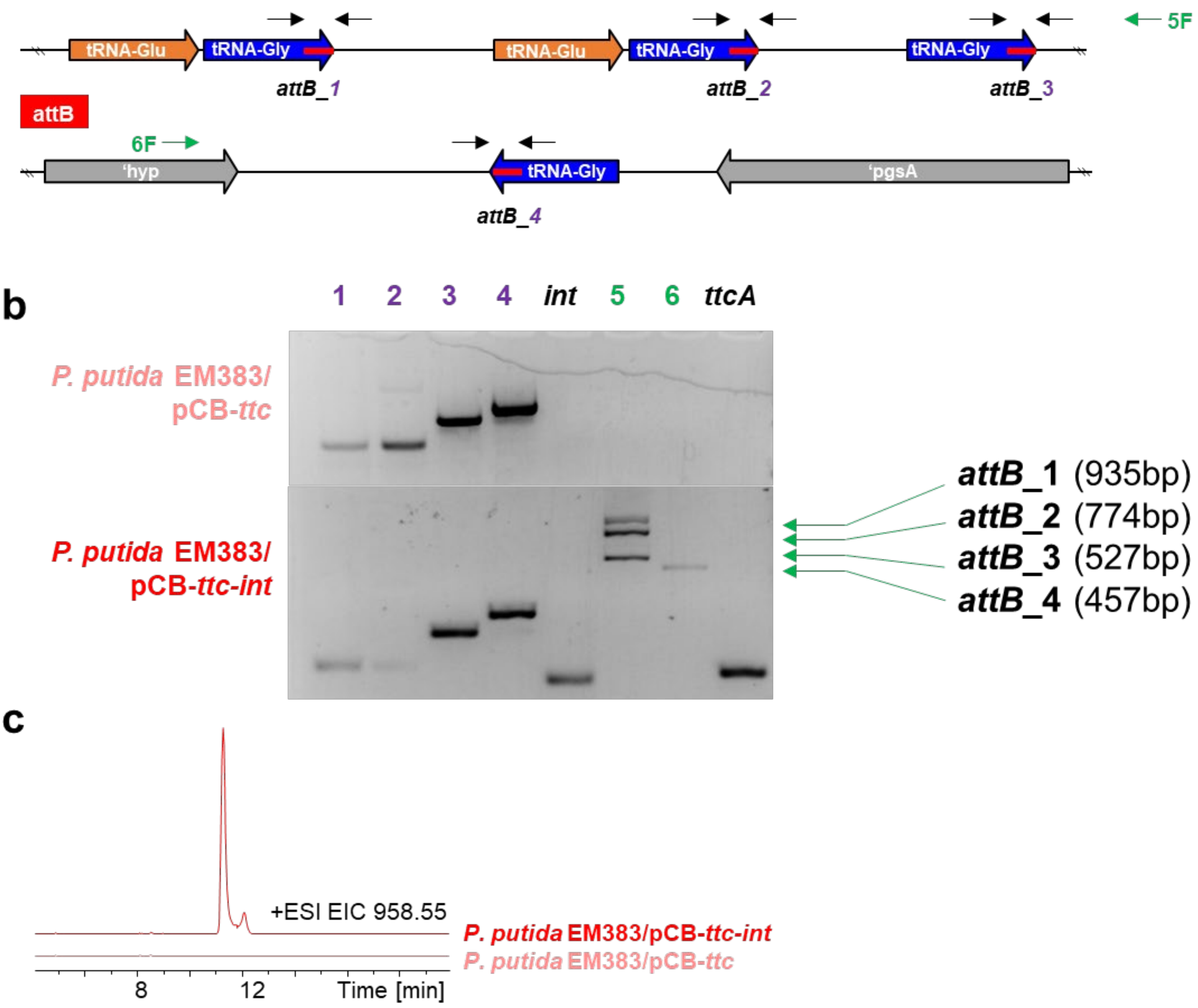


**Supplementary Fig. 2 | Characterization of IntB13-catalyzed integration of *ttc* into the genome of *P. putida* EM383**. **a**) Schematic showing IntB13 attachment sites (*attB*, represented by red bars) in *P. putida* genome, which sit at the 3' end of glycine tRNA genes. Sites 1-3 are clustered within a single locus, while site 4 is distantly located. Attachment sites were amplified across individually using primer pairs 1-4. Alternatively, primers 5F and 6F were paired with a reverse primer that anneals within *intB13* to determine heterogenous integration into multiple attachment sites, or if integration within a single clone was not "clean". Integration into attachment sites 1-3 can be distinguished based on the size of the amplified PCR product. **b**) PCR analysis of individual clones of *P. putida* EM383 into which pCB-*ttc* with and without *intB13* was transferred. Kanamycin resistant *P. putida* EM383 clones can be obtained even in the absence of *intB13*, but *ttc* is not integrated into the genome or stably maintained based on this gel, as the *ttcA* gene cannot be amplified from genomic DNA. Conversely, *intB13* enables stable maintenance, as *ttcA* can be reliably amplified from genomic DNA of clones given pCB-*ttc-int*. However, the integrated construct exists within multiple attachment sites and does not integrate irreversibly within a single site, as attachment into all sites can be detected from a single clone. Many additional clones were tested with similar results (data not shown). **c**) Extracted ion chromatograms (EIC) of thalassospiramide A (**1**) obtained from LC-MS analysis, demonstrating that thalassospiramides are only produced by *P. putida* harboring pCB-*ttc*-*int*.

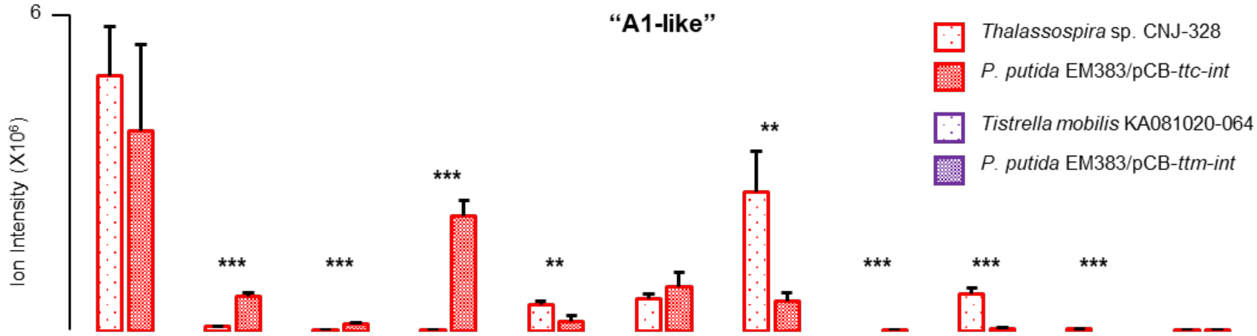


| | A (**1**) | A2 (**2**) | A10 (**3**) | A12 (**4**) | A8 (**5**) | A1 (**10**) | A5 (**8**) | C1 (**11**) | C2 (**6**) | C3* (**9**) | E2 (**7**) |
|---|---|---|---|---|---|---|---|---|---|---|---|
| $AA_1$ | Ser | Ser | Ser | Ser | Ser | Tyr | Phe | Tyr | Ser | Phe | Ser |
| $R_1$ | H | H | H | H | $CH_3$ | H | H | H | H | H | H |
| n | 1 | 1 | 1 | 1 | 1 | 1 | 1 | 0 | 0 | 0 | 2 |
| FA | C10:1 | C10 | C8 | C10-OH | C10:1 | C10:1 | C10:1 | C10:0 | C10:1 | C10:1 | C10:1 |

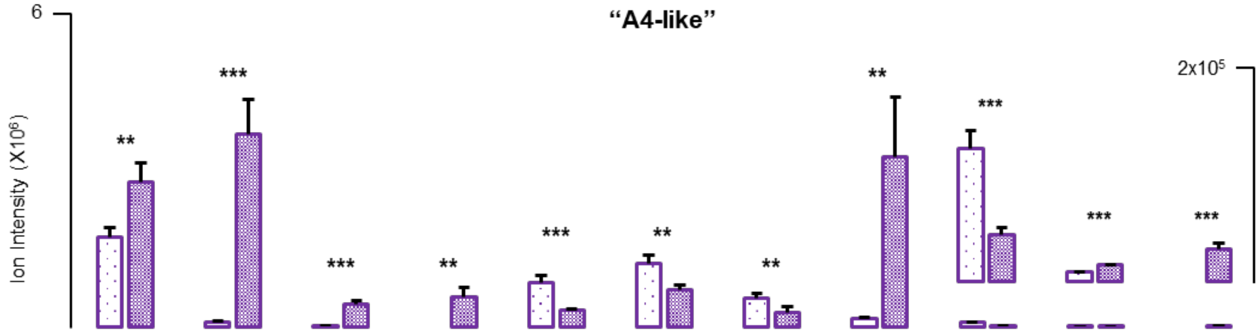


| | A (**1**) | A2 (**2**) | A10 (**3**) | A12 (**4**) | A8 (**5**) | A13 (**19**) | A14 (**20**) | A4 (**21**) | C2 (**6**) | E2 (**7**) | E3 (**22**) |
|---|---|---|---|---|---|---|---|---|---|---|---|
| $AA_1$ | Ser | Ser | Ser | Ser | Ser | Ser | Ser | Val | Ser | Ser | Val |
| $R_1$ | H | H | H | H | $CH_3$ | H | $CH_3$ | H | H | H | H |
| $R_2$ | H | H | H | H | H | $CH_3$ | $CH_3$ | H | H | H | H |
| n | 1 | 1 | 1 | 1 | 1 | 1 | 1 | 1 | 0 | 2 | 2 |
| FA | C10:1 | C10 | C8 | C10-OH | C10:1 | C10:1 | C10:1 | C10:1 | C10:1 | C10:1 | C10:1 |

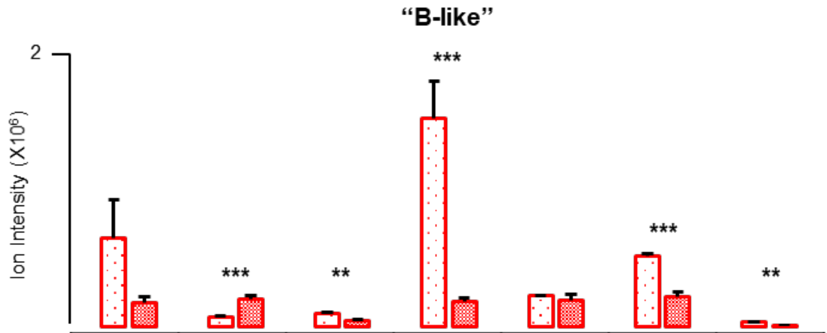


| | B (**12**) | B1 (**15**) | B2 (**17**) | D (**13**) | D1 (**16**) | D2 (**18**) | F (**14**) |
|---|---|---|---|---|---|---|---|
| $AA_1$ | Phe | Tyr | Ser | Phe | Tyr | Ser | Phe |
| n | 1 | 1 | 1 | 0 | 0 | 0 | 2 |

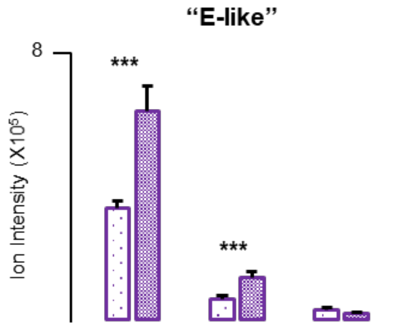


| | E (**23**) | E1 (**24**) | E4 (**25**) |
|---|---|---|---|
| $AA_1$ | Val | Val | Val |
| n | 1 | 2 | 0 |

**Supplementary Fig. 3 | Heterologous production of thalassospiramide analogs from wild-type producers and heterologous host determine by LC-MS.** Extractions were made from triplicate 50 mL cultures and MS ion intensities were normalized by culture density at time of extraction. Significance was determined using a two-tailed Student’s T test, **p<0.05, ***p<0.005. For full structures, see **Fig. 2** in main text. Asterix indicates that *C3 (**9**) was the only analog detected from wild-type but not the host.

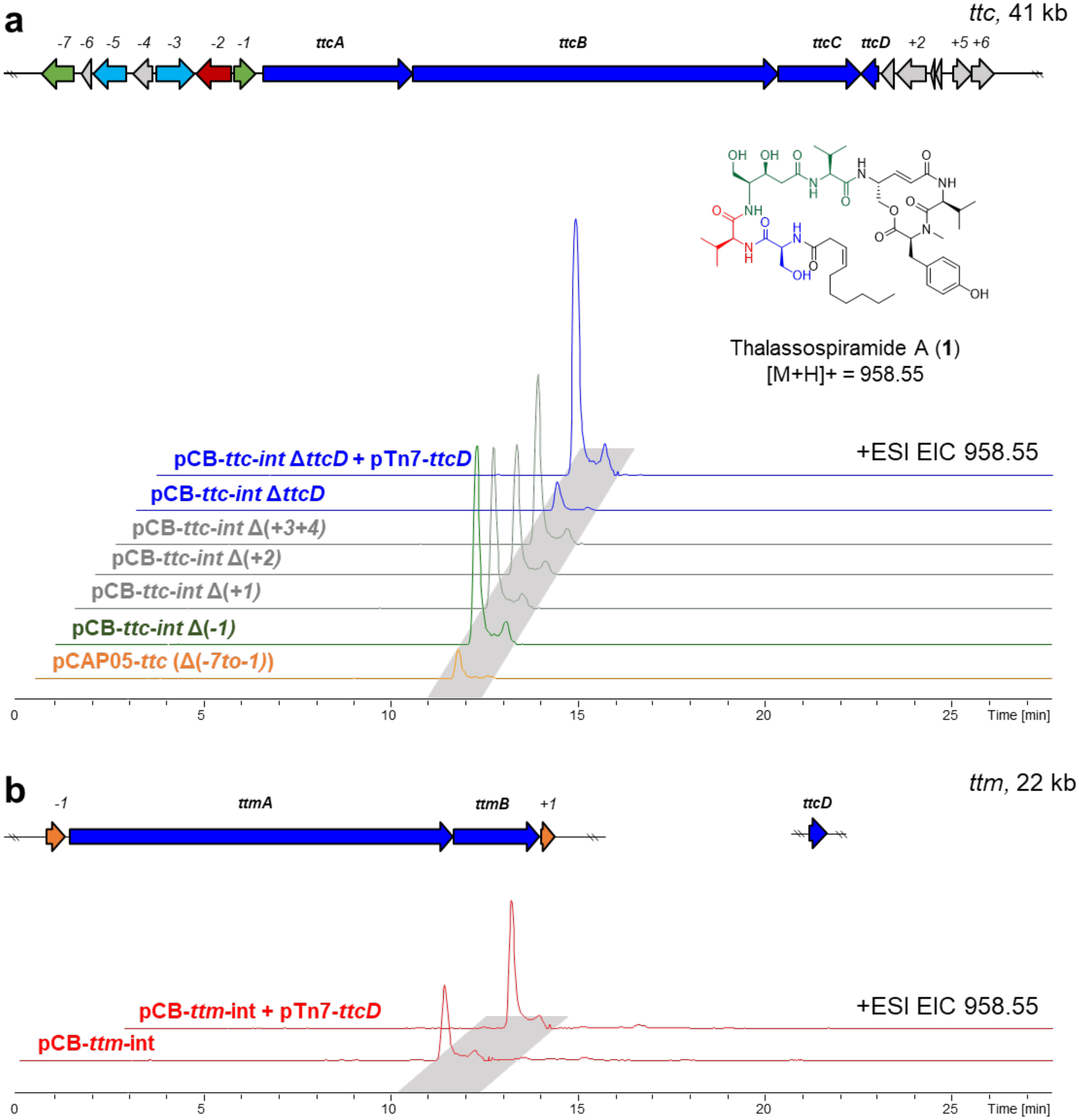


**Supplementary Fig. 4 | Deletion and complementation of non-assembly line genes. a**) EICs for heterologously produced thalassospiramide A (**1**) following deletion or complementation of non-assembly line genes. Heterologous expression of *ttc* using pCAP05 (orange trace) did not include any upstream genes (*-7* through *-1*); although all thalassospiramides, including A (**1**), could be detected, the construct was found to be genetically unstable over time and produce low yields. Individual deletion of upstream (*-1*, green trace) or downstream (*+1* through *+4*, gray traces) genes in the stable pCB-*ttc*-*int* construct did not affect production of thalassospiramide A (**1**), although deletion of *ttcD* resulted in dramatic reduction which could be restored upon genetic complementation (blue traces). **b**) EIC traces showing co-expression of *ttcD* and *ttm* genes increases heterologous thalassospiramide A (**1**) production.

**a**

| *ttc* Experiment 1 | | | | | | | | | | | | | |
|---|---|---|---|---|---|---|---|---|---|---|---|---|---|
| | "A1-like" | | | | | | "B-like" | | | | | | |
| | A (**1**) | A1 (**10**) | A5 (**8**) | C1 (**11**) | C2 (**6**) | E2 (**7**) | B (**12**) | B1 (**15**) | B2 (**17**) | D (**13**) | D1 (**16**) | D2 (**18**) | F (**14**) |
| AA$_1$ | Ser | Tyr | Phe | Tyr | Ser | Ser | Phe | Tyr | Ser | Phe | Tyr | Ser | Phe |
| n | 1 | 1 | 1 | 0 | 0 | 2 | 1 | 1 | 1 | 0 | 0 | 0 | 2 |
| +*ttcD* / Δ*ttcD* | 4.13 ** | 4.59 | 6.33 ** | 2.44 ** | 1.94 * | 4.97 ** | 6.13 ** | 6.67 ** | 5.38 ** | 3.80 ** | 3.72 ** | 3.18 * | 6.09 ** |
| X+/A+ / XΔ/AΔ | | *1.11* | *1.53* | *0.59* | *0.47* | *1.21* | *1.49* | *1.62* | *1.30* | *0.92* | *0.90* | *0.77* | *1.48* |

| *ttc* Experiment 2 | | | | | | | | | | | | | |
|---|---|---|---|---|---|---|---|---|---|---|---|---|---|
| | "A1-like" | | | | | | "B-like" | | | | | | |
| | A (**1**) | A1 (**10**) | A5 (**8**) | C1 (**11**) | C2 (**6**) | E2 (**7**) | B (**12**) | B1 (**15**) | B2 (**17**) | D (**13**) | D1 (**16**) | D2 (**18**) | F (**14**) |
| AA$_1$ | Ser | Tyr | Phe | Tyr | Ser | Ser | Phe | Tyr | Ser | Phe | Tyr | Ser | Phe |
| n | 1 | 1 | 1 | 0 | 0 | 2 | 1 | 1 | 1 | 0 | 0 | 0 | 2 |
| +*ttcD* / Δ*ttcD* | 5.18 * | 9.83 | 12.13 ** | 2.78 * | 3.87 ** | 5.84 * | 8.31 ** | 5.97 * | 4.48 ** | 3.77 ** | 2.80 * | 2.35 * | 5.19 ** |
| X+/A+ / XΔ/AΔ | | *1.90* | *2.34* | *0.54* | *0.75* | *1.13* | *1.60* | *1.15* | *0.86* | *0.73* | *0.54* | *0.45* | *1.00* |

**b**

| *ttm* Experiment 1 | | | | | | | | |
|---|---|---|---|---|---|---|---|---|
| | "A4-like" | | | | | "E-like" | | |
| | A (**1**) | A4 (**21**) | C2 (**6**) | E2 (**7**) | E3 (**22**) | E (**23**) | E1 (**24**) | E4 (**25**) |
| AA$_1$ | Ser | Val | Ser | Ser | Val | Val | Val | Val |
| n | 1 | 1 | 0 | 2 | 2 | 1 | 2 | 0 |
| +*ttcD* / -*ttcD* | 1.51 * | 2.08 * | 0.60 * | 10.3 * | 12.7 * | 1.09 | 4.13 ** | 0.68 |
| X+/A+ / X–/A– | | 1.38 | 0.40 | 6.81 | 8.44 | 0.72 | 2.75 | 0.45 |

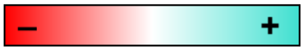


| *ttm* Experiment 2 | | | | | | | | |
|---|---|---|---|---|---|---|---|---|
| | "A4-like" | | | | | "E-like" | | |
| | A (**1**) | A4 (**21**) | C2 (**6**) | E2 (**7**) | E3 (**22**) | E (**23**) | E1 (**24**) | E4 (**25**) |
| AA$_1$ | Ser | Val | Ser | Ser | Val | Val | Val | Val |
| n | 1 | 1 | 0 | 2 | 2 | 1 | 2 | 0 |
| +*ttcD* / -*ttcD* | 2.61 ** | 2.56 * | 0.57 ** | 9.7 * | 10.4 ** | 1.2 | 3.61 ** | 0.70 |
| X+/A+ / X–/A– | | *0.98* | *0.34* | *5.67* | *6.07* | *0.74* | *2.11* | *0.41* |

**Supplementary Fig. 5 l Quantitative analysis of TtcD effect on thalassospiramide product distribution. a**) Results of two independent experiments comparing host cultures expressing *ttc* with *ttcD* deleted or re-complemented. Extractions were made from triplicate 50 mL cultures and MS ion intensities were normalized by culture density at time of extraction. The top row of values represents the fold-change in ion intensity for a given analog upon *ttcD* complementation. For the bottom row, ion intensities have been normalized to thalassospiramide A (**1**) levels from the same culture and then compared to represent normalized fold-change value. Color intensity corresponds to numerical value. Significance was determined using a two-tailed Student's T test, *$p<0.05$, **$p<0.005$. For full chemical structures, see **Fig. 2** of main text. As we hypothesize the repeat ser-C2-val units arise from pass-back chain extension, these results suggest that TtcD promotes this activity to favor formation of analogs in which n≥1. **b**) Results of two independent experiments comparing host cultures expressing *ttm* with or without *ttcD* co-expression. Co-expression favorably enhances production of analogs with n≥1.

**a**

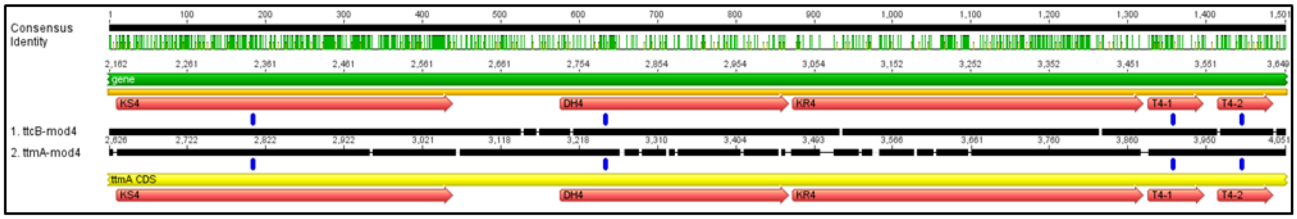

**b**

```
Ery-DH4         ----------DVSALGV----RGAEHPLLLAAVDVPGHGGAVFTGRLSTDEQPWLAEHVV   46
ttcB-DH4        LPHAPFAKIVCKPVFGS---NRAPDIAIMTGPVE--TERGRCFAIPVGTPSFWPTGEHHL   2789
ttmA-DH4        ----PFSRIPCRIDLPAKAANSDIDTTFSMGWTA--FPGGRALALPLADPAFWPVAEHRL   3253
                              :         :  :  . .      *  ::  :.       .** :

Ery-DH4         GGRTLVPGSVLVDL---ALAAGEDVGLPVLEELVLQRPLVLAG----AGALLRMSVGAPD   99
ttcB-DH4        NGQPTLVGMAVPAMIAAAIRNGTGDPAVRISDLKWQKTLLPNALPDGTATLLLGNDGLV-   2848
ttmA-DH4        AGRPTLVGMAIPALVAADR-----GPGTVLHDLVWRRPLHAG---SREASLVIARDGSV-   3304
                 *:  : * .:  :               : :*  :: *         .:*:    *

Ery-DH4         ESGRRTIDVHAAEDVADLADAQWSQHATGTLAQGVAAGPRDTEQWPPEDAVR----IPLD   155
ttcB-DH4        EL------------GGRLKNGKWSVFATARWANDAPESSH-HDNLPSLDHARAKCTLAAN   2895
ttmA-DH4        TA------------GHATAEGGWAVAAEARVETSSAP-------HPSAVDLSTDGLIALD   3345
                                   :. *:  * .    .           *          :  :

Ery-DH4         -DHYDGLAEQGYEYGPSFQALRAAWRKDDSVYA------EVSIAADEEGYAFHPVLLDAV   208
ttcB-DH4        IAPYAP-EIGPITISDRWDCRVSMQYSADETMAMMHLKLKPQYHADLRDWVIHPAMADIA   2954
ttmA-DH4        LAPFDG-AEGVMQVGPRWDCRQAIWLAPDRRRAVARLALPAAAAADRRLWPWHPALLDIA   3404
                   :          .  ::.  :     *   *           ** . :  **.: * .

Ery-DH4         AQTLSLGALGEPGGGKLPFAWNTVTLHASGATSVRVVATPAGADAMALRVTDPAGHL-VA   267
ttcB-DH4        CSMIL----TAEDNGSIPVGVDEITLHAPFTDDILVCTERHKPGQADFSFFDAQSQKLLL   3010
ttmA-DH4        ASLLA-------GVGEVPREVAAIRILAPLPDTVMARADRRPDGLVDIRLADAAGRP-CL   3457
                .. :        . *.:*     : : *     : . :     .   : . *  .:

Ery-DH4         TVDSLVVR-----   275
ttcB-DH4        TIRGIRFSR----   3019
ttmA-DH4        LIDGLRFVAADGR   3469
                 : .: .
```

**Supplementary Fig. 6 | Annotation and alignment of DH₄.** **a**) Annotation of unusual dehydratase domain of module 4 ($DH_4$), which spans approximately 250 amino acids within TtcB and TtmA. **b**) Protein sequence alignment of TtcB and TtmA $DH_4$ with erythromycin (Ery) module 4 dehydratase, generated using Clustal Omega under default settings. Conserved motifs highlighted in yellow and supporting active site residues bolded in Ery-$DH_4$ sequence. Catalytic histidine (H) and aspartic acid (D) residues shown in bold and colored red.

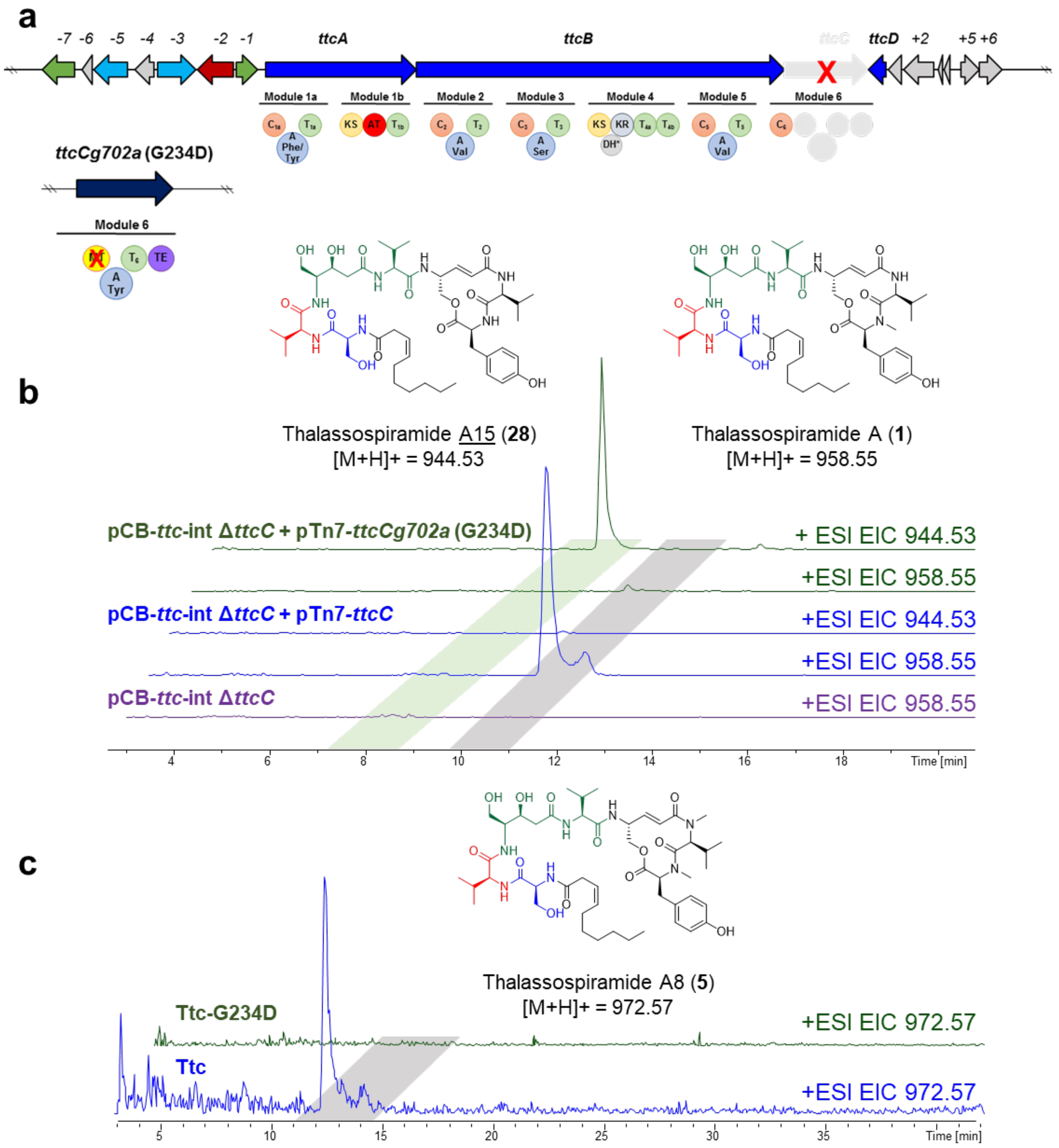


**Supplementary Fig. 7 | Inactivation of $MT_6$ via gene deletion and complementation of *ttcC*. a**) Schematic for gene deletion and complementation strategy for precise inactivation of $MT_6$ encoded on TtcC. **b**) EICs for thalassospiramide A (**1**) and A15 (**28**) from deletion and complementation mutants, showing that $MT_6$ inactivation results in formation of the desmethyl analog. **c**) EIC showing $MT_6$ inactivation results in complete loss of dimethyl analog thalassospiramide A8 (**5**).

**C domains**

```
TtcA-C1a         VLIAHNEVEEIRRRPLDPENGRHCRHRLLQLGKGRFWWVRVYHHLVCDGYAGHLMAMRAA    170
TtmA-C1          DLLAQEAFEALQARPFDTAQGPLCRHQLLRLAPDRHRWIRCYHHLILDGQGGMILAGAVG    179
TtmA-C2          IRLAGYDDAAEMLAPFDPATGPLIRFGWEADQPERLR--VVVDHLVFDGESRTVFQR---    713
                   :*          *:*   *   *.        *       .**: ** .  ::
```

**A domains (core A3)**

```
GrsA-A           EDTIKIREGT-NL---HVPSKSTDLAYVIYTSGTTGNPKGTMLEHKGISNLKVFFENSLN   219
TtcA-A1a         INAVPGIDGRTVLEDVLADICGDDIAFVFHTSGSTGQPKPVPVHHQSLADKIDVAIAQFG   658
TtcB-A3          ANNVLIIDGSEDADIPDAINAPDDLAYVLFTSGSTGRPKGVEIAHRGVLNRILWMQDAFP   1733
                  : :   :*        .     *:*:*:.***:**.** . : *:.: :        :
```

**A domains (core A10)**

```
GrsA-A           LPTYMIPSYFIQLDKMPLTSNGKIDRKQLPEPDLTFGMRV--DYEAPRNEIEETLVTIWQ   552
TtcA-A1a         LSDAAVPTRLEWVETLPLLPSGKIDRKALAQFAQAPGGKQPVAQETPPPASRKPDQMRIH   1009
TtcB-A3          LPEYMIPARFFALDHLPLTSSGKVDRKALSGTPMAGSPKT--ASRKSKPAIAAKPDAEIH   2062
                 *    :*: :  :: :**  .**:*** *     : . :     .              :
```

**T domains**

```
TtcA-T1a         AEKIAKIWMDLIETDEIEFDTNLFEAGAHSLLVPRAQFALSKLAGRNIASVEIFQHPTIN   1071
TtcB-T4a         PQRIAAIWAETLGYDAVASDDDFFALGGDSITGMQIVDRINAELKLSLAISDLFAAPTVS   3539
TtcB-T4b         -DRIAKIWAEILGYDAIDPDEDFYALGGDSITGMQIVDRMNAELACNIGLADLFETPTIT   3625
                  ::** ** : :  * :  * :::  *..*:   :    :.     .:.  ::*  **:.
```

**KS domains**

```
DEBS-KS2          VGLIPQEY-----GPRLAEGGEGVEGYLMTGTTTSVASGRIAYTLGLEGPAISVDTACSS  732
TtcA-KS1b         VGVGFPTYLVDSLRDRLRDRPDAIRYGMTLGNDKDFAATRLAYKLNLTGPAVASSTACST  1276
                  **:    *       ** :  :.:.  :  *. ...*: *:**.*.* ***:: .****:
```

**AT domains**

```
DEBS-AT1          SRRVEVVQPALFAVQTSLAALWRSFGVTPDAVVGHSIGELAAAHVCGAAGAADAARAAAL  221
TtcA-AT1b         MARTEIAQPALFIYEYALASWLIGNGISPAALAGHSIGEYVAACIGGVMSFDDALRLVVT  1752
                    *.*:.*****  : :**:   . *::* *:.****** .** : *. .  ** * ..
```

**MT domains**

```
AcmC-MT4         GLPIPLDQMREWRDTTVERIRGLNP--RRVLEIGVGTGLLLSRLAPHCEEYWGTDFSPTV   2073
TtcC-MT6         GQPIAQSAMDAFGANARNKVAGLVSRDARVLEIGCASGFTMKHVAPVVGTYVATDLSRRN   260
TtmB-MT6         GQPIAEAVMAEFGQAAAGKLSGLVTPASRVLEIGCASGFTLRALAPLAGPYLATDISRRA   256
                 * **    *  :   :  :: **     ****** .:*: :  :**    * .**:*
```

**Supplementary Fig. 8 | Protein sequence alignments for identification of Ttc and Ttm domain active sites targeted for mutation.** Alignments generated using Clustal Omega with default settings. Sequence motifs highlighted in yellow; conserved residues bolded and active sites targeted for mutation colored in red. C, condensation; A, adenylation; T, thiolation; KS, ketosynthase; AT, acyltransferase; MT, methyltransferase; Grs, gramicidin synthetase; DEBS, 6-deoxyerythronolide B synthase; Acm, actinomycin synthetase.

**a**

| | Name | | Name |
|---|---|---|---|
| 1 | pCB-*ttc-int* | 10 | pCB-*ttc-int*-A3 |
| 2 | pCB-*ttc-int*-C1a 1 | 11 | pCB-*ttc-int*-T4 A |
| 3 | pCB-*ttc-int*-C1a 2 | 12 | pCB-*ttc-int*-T4 B |
| 4 | pCB-*ttc-int*-A1a (G631D) | 13 | pCB-*ttm-int* |
| 5 | pCB-*ttc-int*-A1a (K972A) 1 | 14 | pCB-*ttm-int*-C1 1 |
| 6 | pCB-*ttc-int*-A1a (K972A) 2 | 15 | pCB-*ttm-int*-C1 2 |
| 7 | pCB-*ttc-int*-T1a | 16 | pCB-*ttm-int*-C2 1 |
| 8 | pCB-*ttc-int*-KS1b | 17 | pCB-*ttm-int*-C2 2 |
| 9 | pCB-*ttc-int*-AT1b | 18 | pCB-*ttm-int*-MT6 |

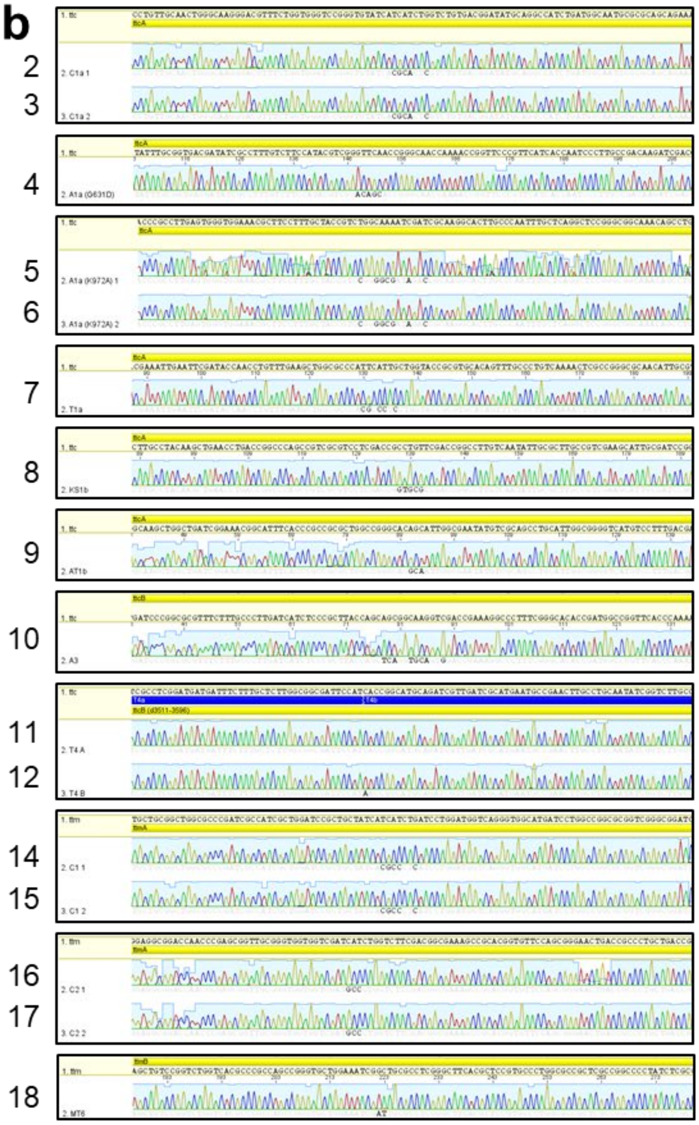


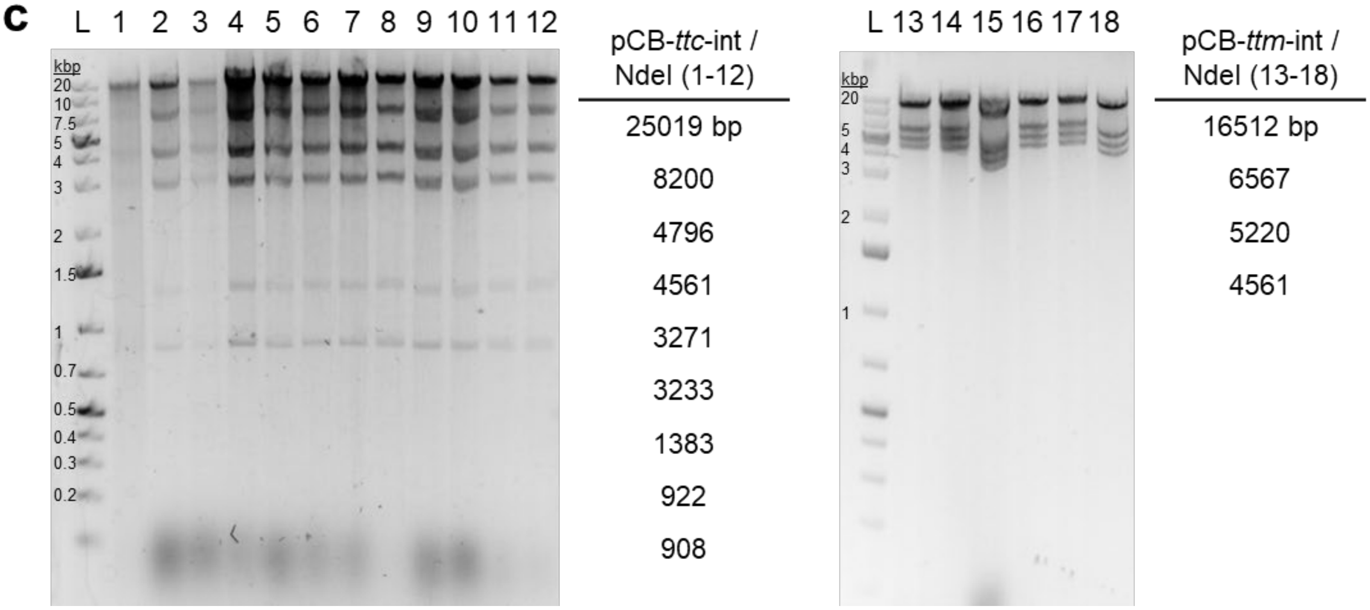


**Supplementary Fig. 9 | Verification of edited constructs. a**) List of constructs generated. **b**) Sanger sequencing results confirming mutations. **c**) Restriction digestion verification.

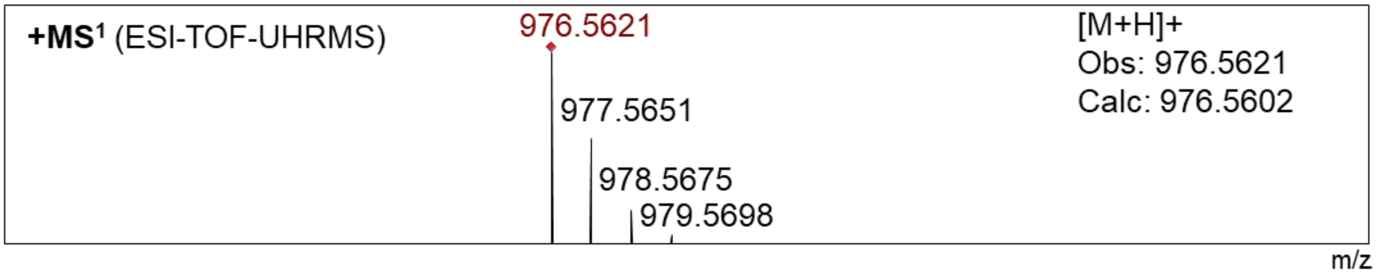


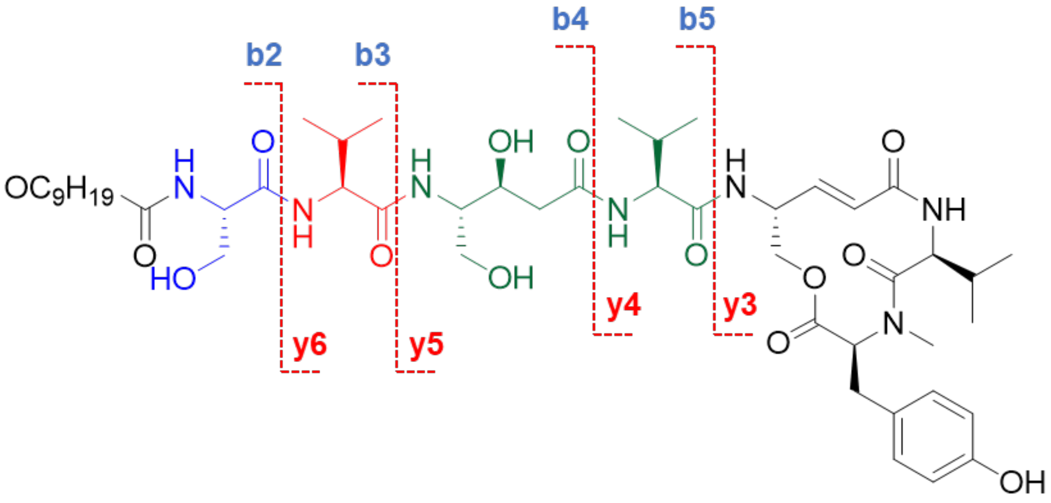


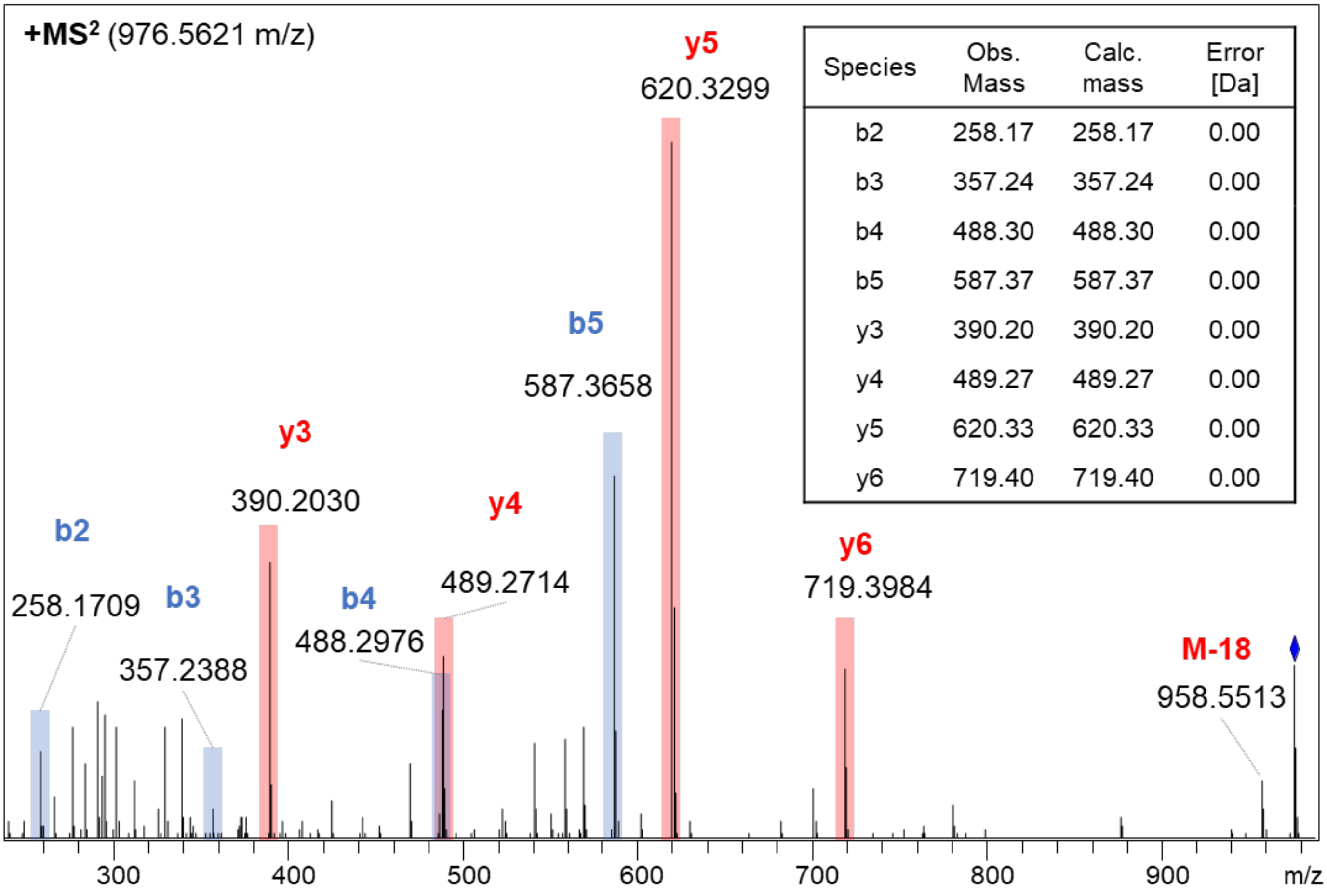


| Species | Obs. Mass | Calc. mass | Error [Da] |
|---|---|---|---|
| b2 | 258.17 | 258.17 | 0.00 |
| b3 | 357.24 | 357.24 | 0.00 |
| b4 | 488.30 | 488.30 | 0.00 |
| b5 | 587.37 | 587.37 | 0.00 |
| y3 | 390.20 | 390.20 | 0.00 |
| y4 | 489.27 | 489.27 | 0.00 |
| y5 | 620.33 | 620.33 | 0.00 |
| y6 | 719.40 | 719.40 | 0.00 |

**Supplementary Fig. 10 | Characterization of thalassospiramide A12 (4).** MS$^n$ analysis.

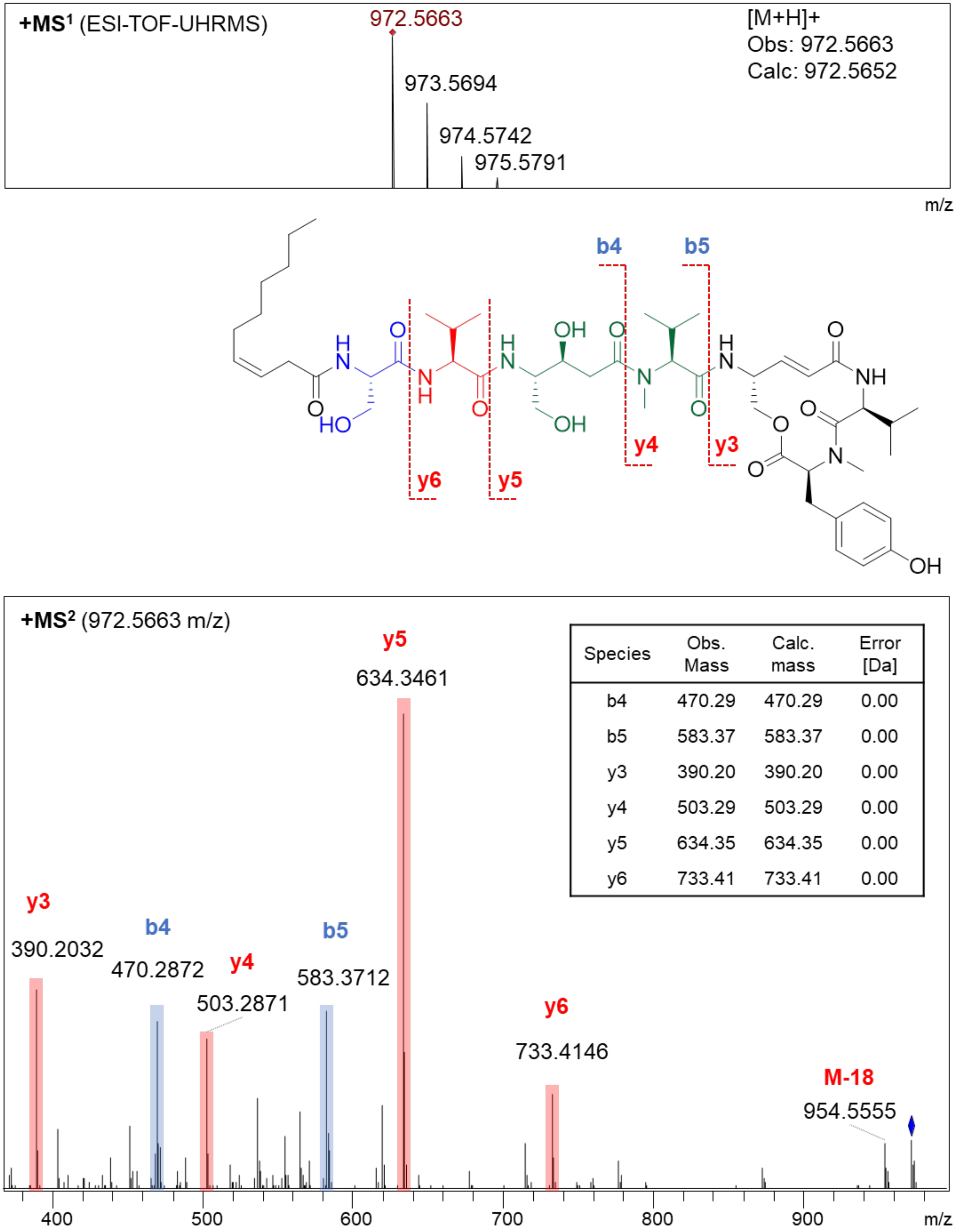


| Species | Obs. Mass | Calc. mass | Error [Da] |
|---|---|---|---|
| b4 | 470.29 | 470.29 | 0.00 |
| b5 | 583.37 | 583.37 | 0.00 |
| y3 | 390.20 | 390.20 | 0.00 |
| y4 | 503.29 | 503.29 | 0.00 |
| y5 | 634.35 | 634.35 | 0.00 |
| y6 | 733.41 | 733.41 | 0.00 |

**Supplementary Fig. 11 | Characterization of thalassospiramide A13 (19).** $MS^n$ analysis.

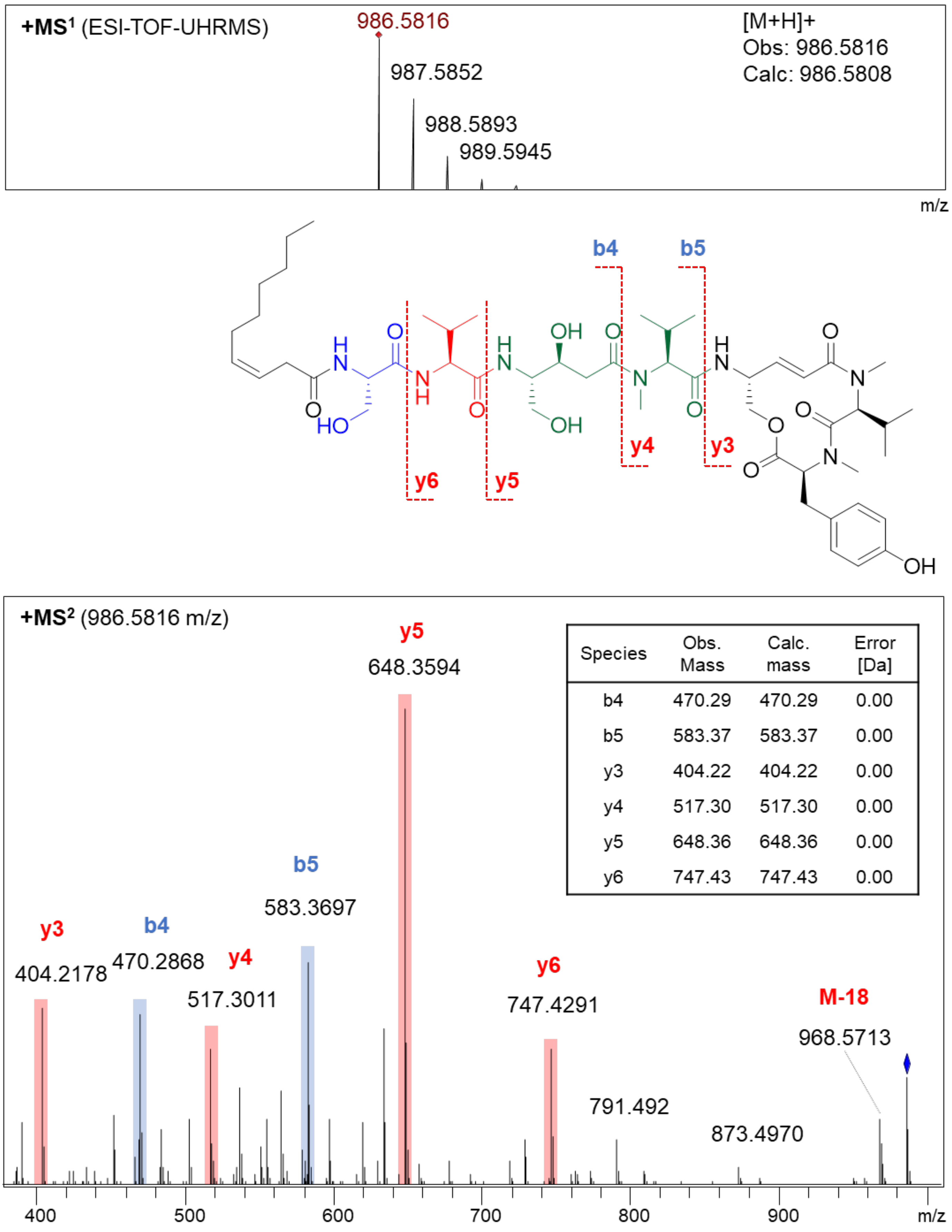


| Species | Obs. Mass | Calc. mass | Error [Da] |
|---|---|---|---|
| b4 | 470.29 | 470.29 | 0.00 |
| b5 | 583.37 | 583.37 | 0.00 |
| y3 | 404.22 | 404.22 | 0.00 |
| y4 | 517.30 | 517.30 | 0.00 |
| y5 | 648.36 | 648.36 | 0.00 |
| y6 | 747.43 | 747.43 | 0.00 |

**Supplementary Fig. 12 | Characterization of thalassospiramide A14 (20).** MS$^n$ analysis.

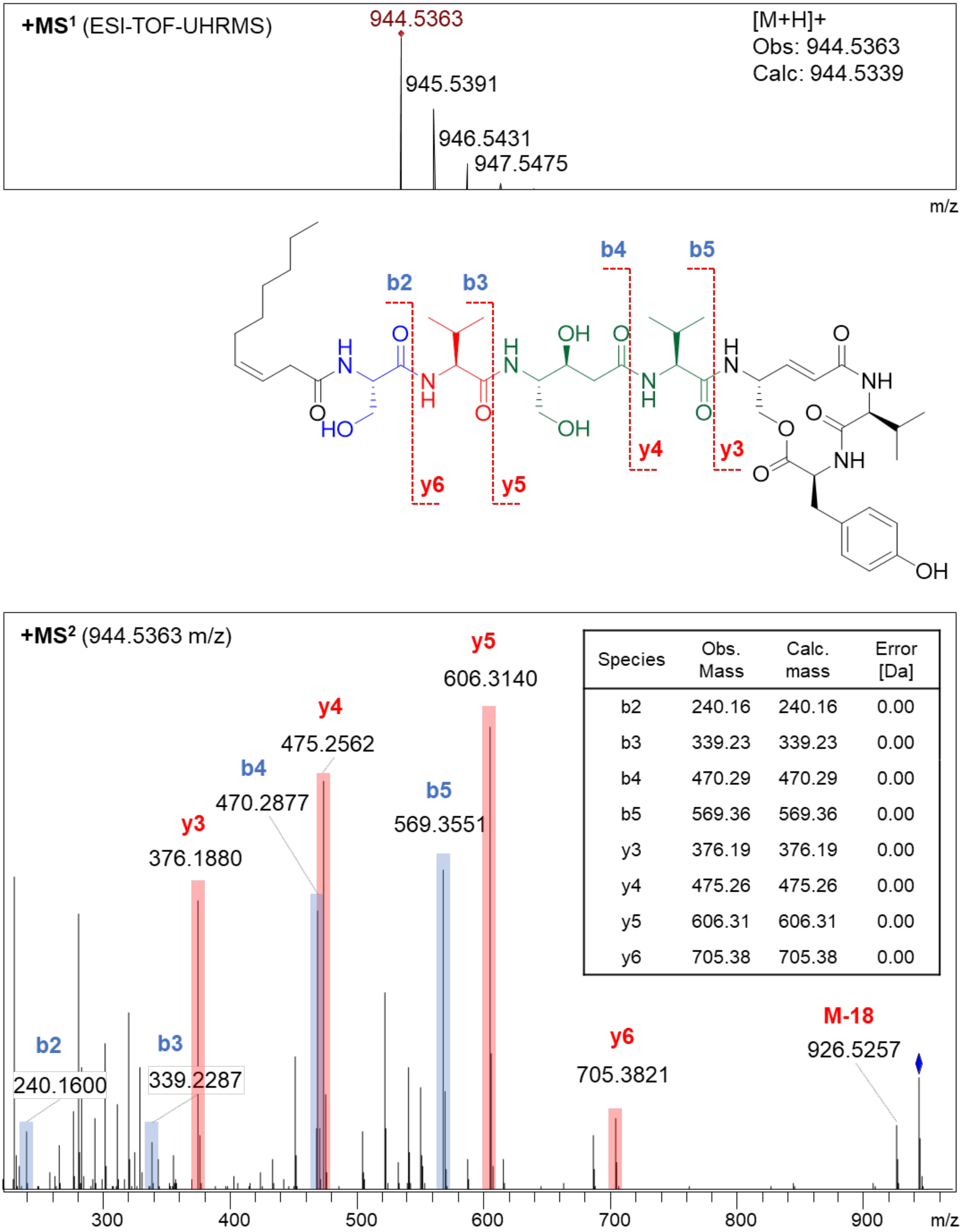


| Species | Obs. Mass | Calc. mass | Error [Da] |
|---|---|---|---|
| b2 | 240.16 | 240.16 | 0.00 |
| b3 | 339.23 | 339.23 | 0.00 |
| b4 | 470.29 | 470.29 | 0.00 |
| b5 | 569.36 | 569.36 | 0.00 |
| y3 | 376.19 | 376.19 | 0.00 |
| y4 | 475.26 | 475.26 | 0.00 |
| y5 | 606.31 | 606.31 | 0.00 |
| y6 | 705.38 | 705.38 | 0.00 |

**Supplementary Fig. 13 | Characterization of thalassospiramide A15 (28).** MS$^n$ analysis.

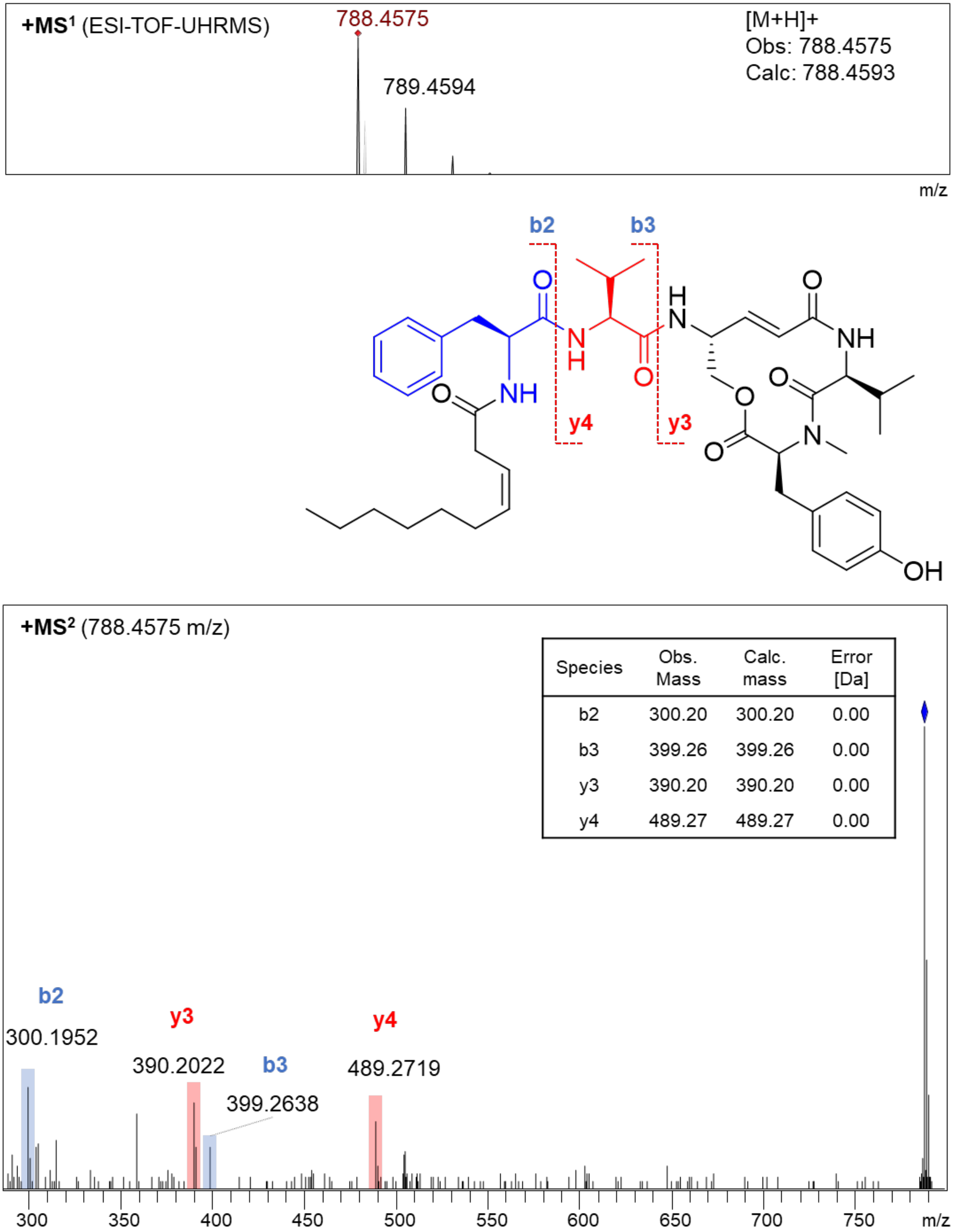


| Species | Obs. Mass | Calc. mass | Error [Da] |
|---|---|---|---|
| b2 | 300.20 | 300.20 | 0.00 |
| b3 | 399.26 | 399.26 | 0.00 |
| y3 | 390.20 | 390.20 | 0.00 |
| y4 | 489.27 | 489.27 | 0.00 |

**Supplementary Fig. 14 | Characterization of thalassospiramide C3 (9).** MSⁿ analysis.

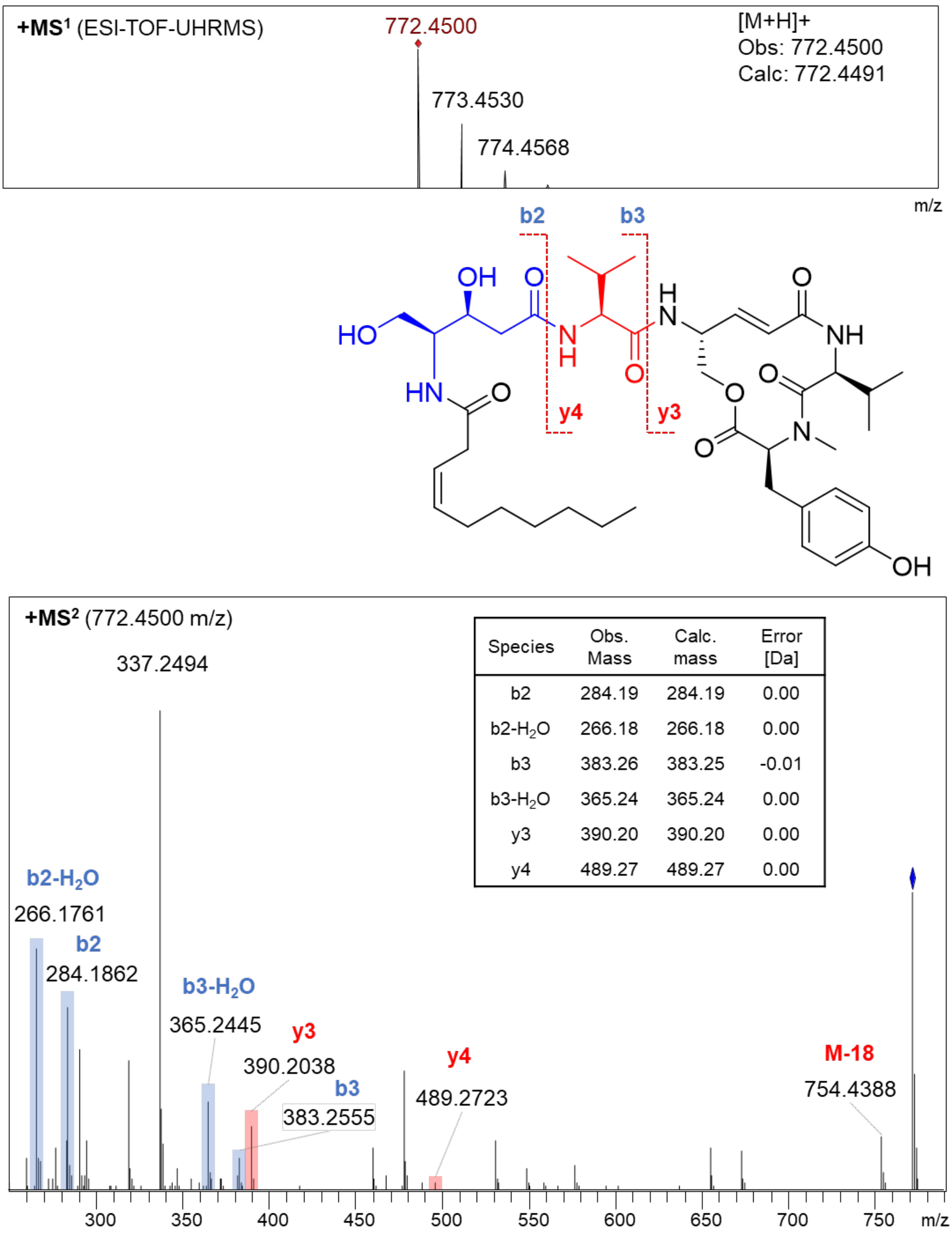


| Species | Obs. Mass | Calc. mass | Error [Da] |
|---|---|---|---|
| b2 | 284.19 | 284.19 | 0.00 |
| b2-$H_2O$ | 266.18 | 266.18 | 0.00 |
| b3 | 383.26 | 383.25 | -0.01 |
| b3-$H_2O$ | 365.24 | 365.24 | 0.00 |
| y3 | 390.20 | 390.20 | 0.00 |
| y4 | 489.27 | 489.27 | 0.00 |

**Supplementary Fig. 15 | Characterization of thalassospiramide D2 (18).** $MS^n$ analysis.

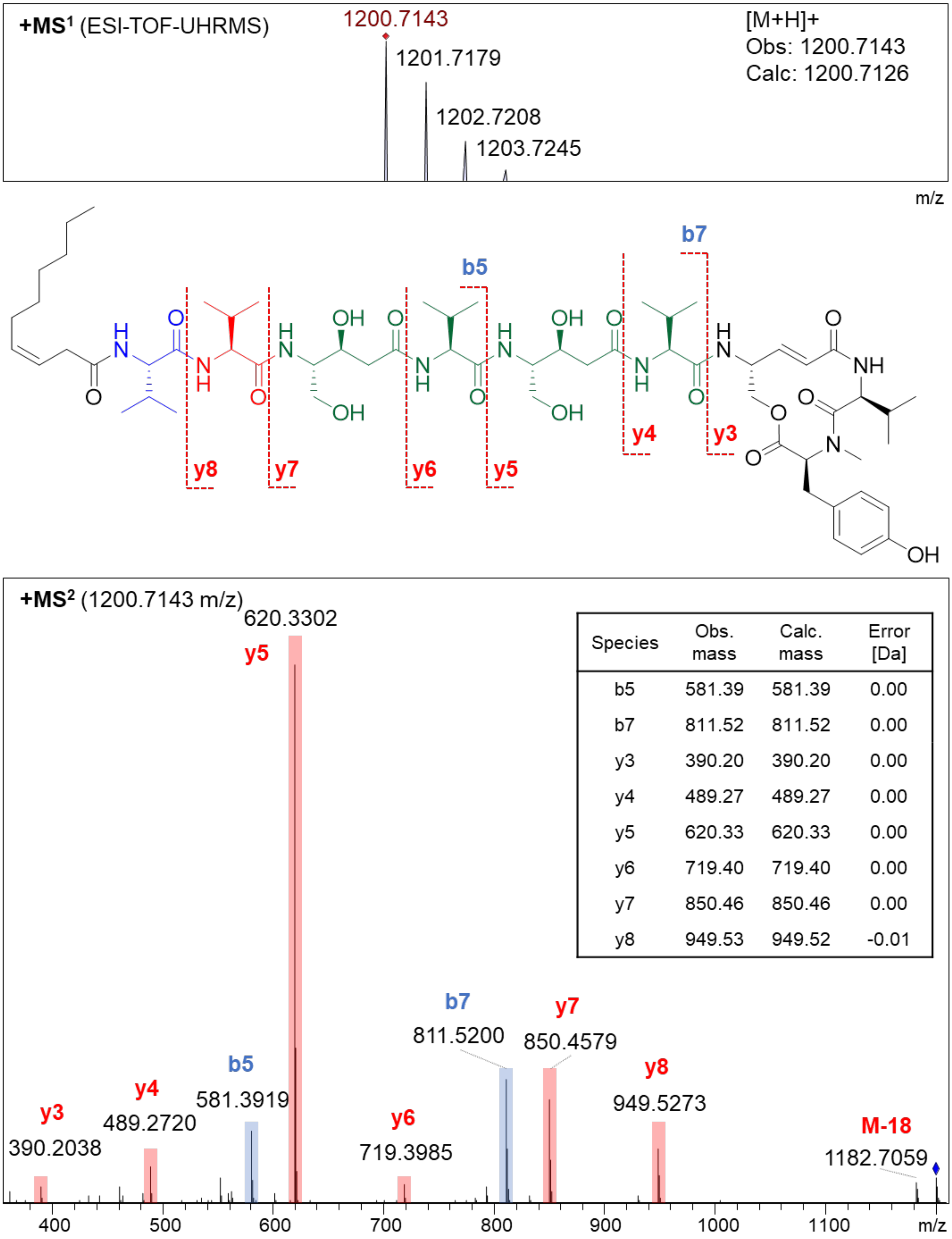


| Species | Obs. mass | Calc. mass | Error [Da] |
|---|---|---|---|
| b5 | 581.39 | 581.39 | 0.00 |
| b7 | 811.52 | 811.52 | 0.00 |
| y3 | 390.20 | 390.20 | 0.00 |
| y4 | 489.27 | 489.27 | 0.00 |
| y5 | 620.33 | 620.33 | 0.00 |
| y6 | 719.40 | 719.40 | 0.00 |
| y7 | 850.46 | 850.46 | 0.00 |
| y8 | 949.53 | 949.52 | -0.01 |

**Supplementary Fig. 16 | Characterization of thalassospiramide E3 (22).** $MS^n$ analysis.

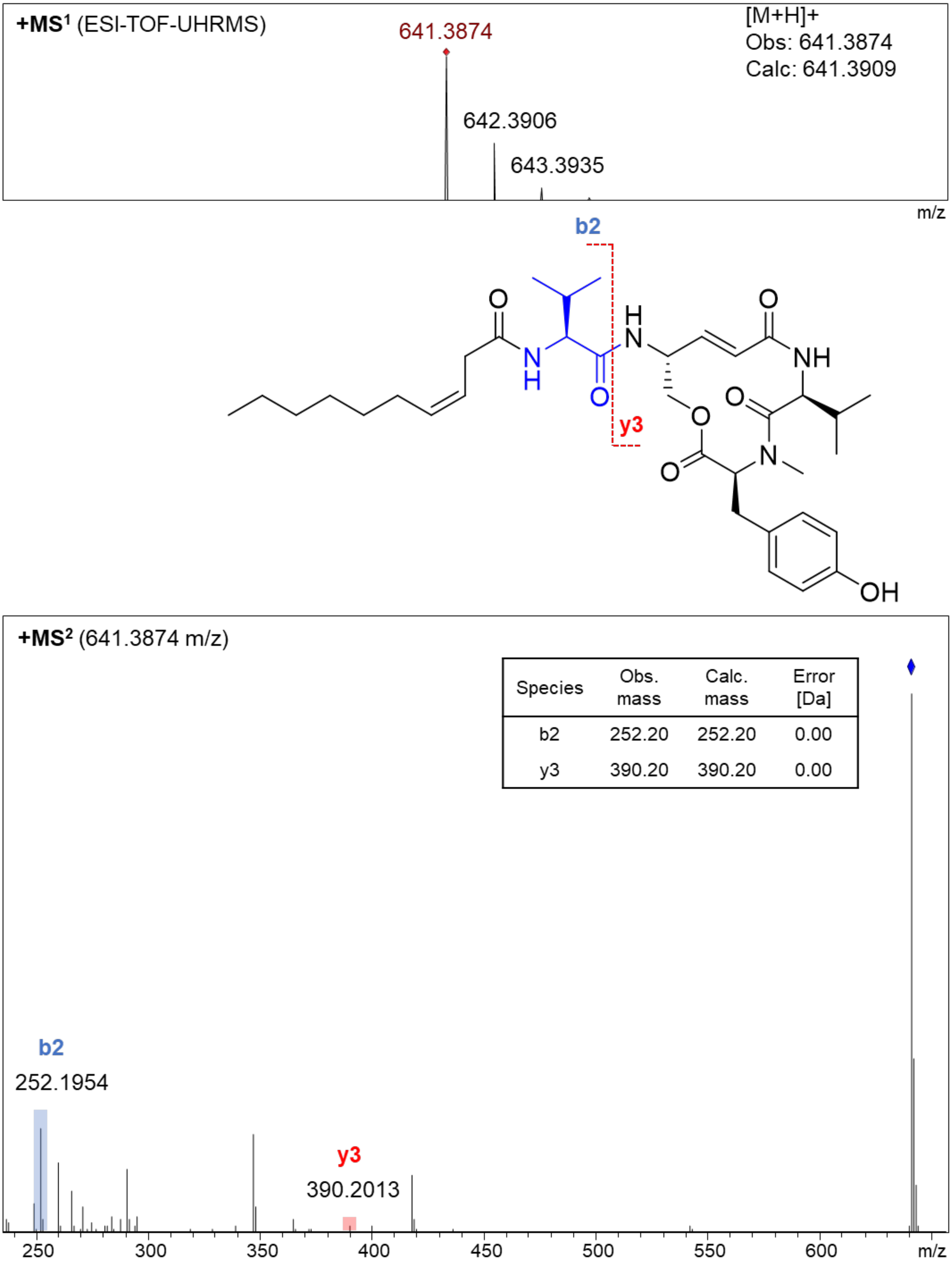


| Species | Obs. mass | Calc. mass | Error [Da] |
|---|---|---|---|
| b2 | 252.20 | 252.20 | 0.00 |
| y3 | 390.20 | 390.20 | 0.00 |

**Supplementary Fig. 17 | Characterization of thalassospiramide E4 (25).** MS$^n$ analysis.

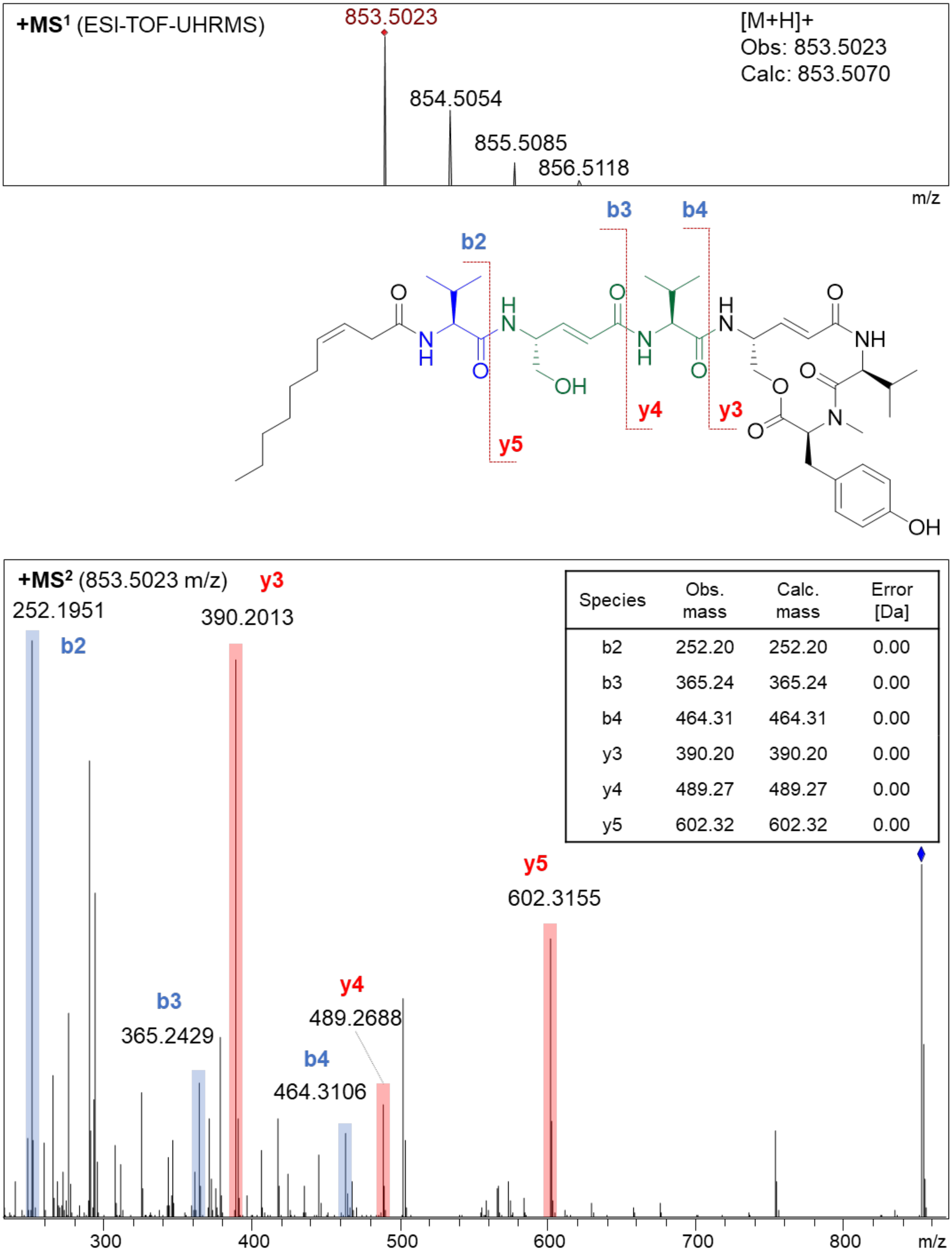


| Species | Obs. mass | Calc. mass | Error [Da] |
|---|---|---|---|
| b2 | 252.20 | 252.20 | 0.00 |
| b3 | 365.24 | 365.24 | 0.00 |
| b4 | 464.31 | 464.31 | 0.00 |
| y3 | 390.20 | 390.20 | 0.00 |
| y4 | 489.27 | 489.27 | 0.00 |
| y5 | 602.32 | 602.32 | 0.00 |

**Supplementary Fig. 18 | Characterization of thalassospiramide <u>E5</u> (26).** MS$^n$ analysis.

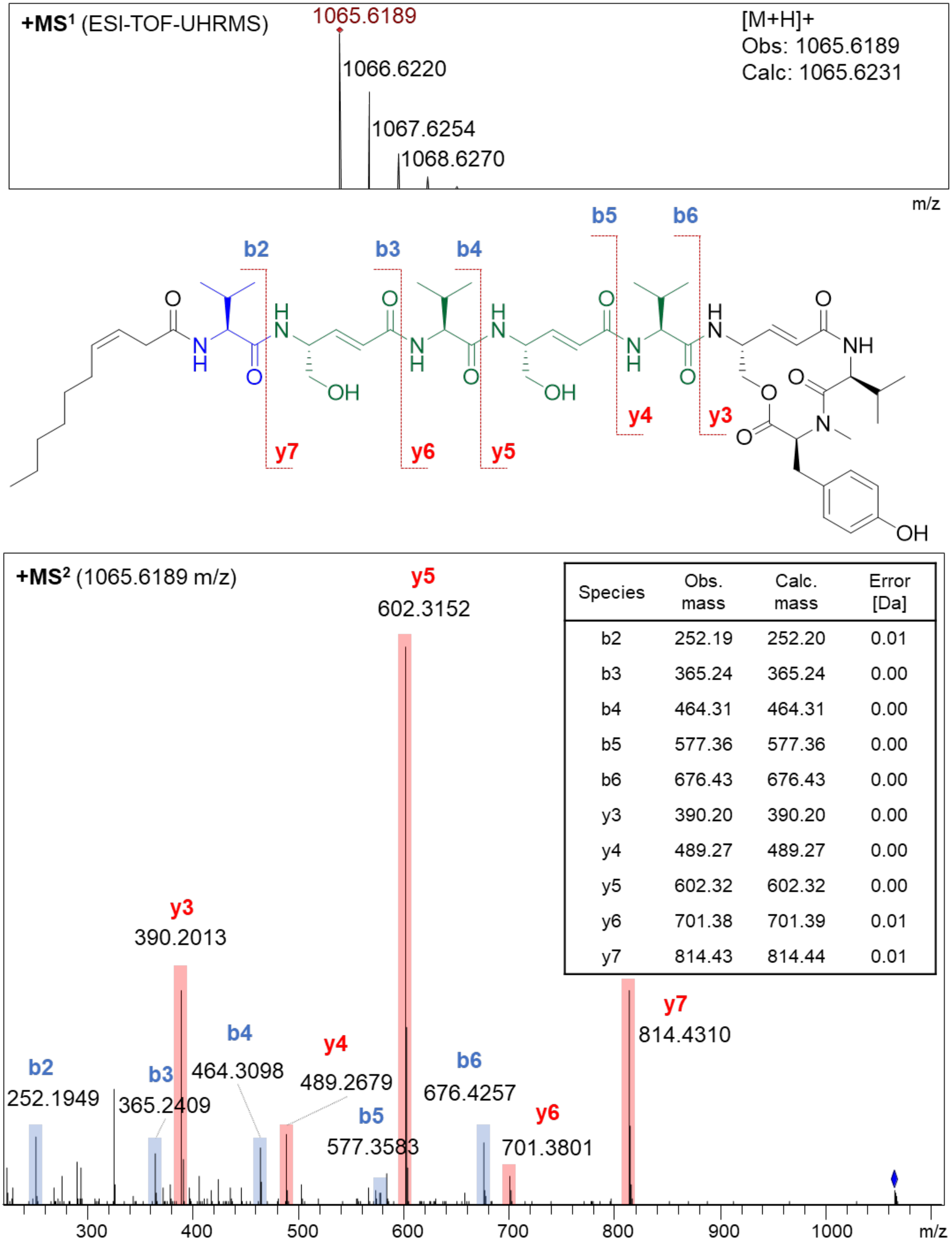


| Species | Obs. mass | Calc. mass | Error [Da] |
|---|---|---|---|
| b2 | 252.19 | 252.20 | 0.01 |
| b3 | 365.24 | 365.24 | 0.00 |
| b4 | 464.31 | 464.31 | 0.00 |
| b5 | 577.36 | 577.36 | 0.00 |
| b6 | 676.43 | 676.43 | 0.00 |
| y3 | 390.20 | 390.20 | 0.00 |
| y4 | 489.27 | 489.27 | 0.00 |
| y5 | 602.32 | 602.32 | 0.00 |
| y6 | 701.38 | 701.39 | 0.01 |
| y7 | 814.43 | 814.44 | 0.01 |

**Supplementary Fig. 19 | Characterization of thalassospiramide E6 (27).** $MS^n$ analysis.

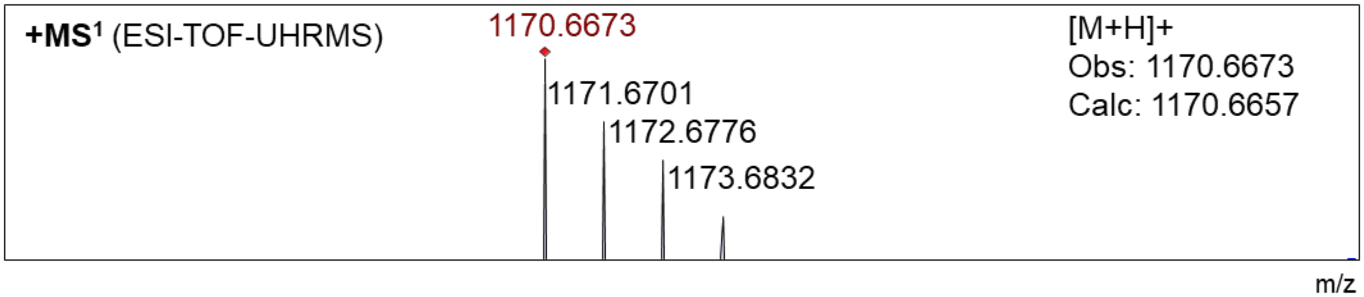


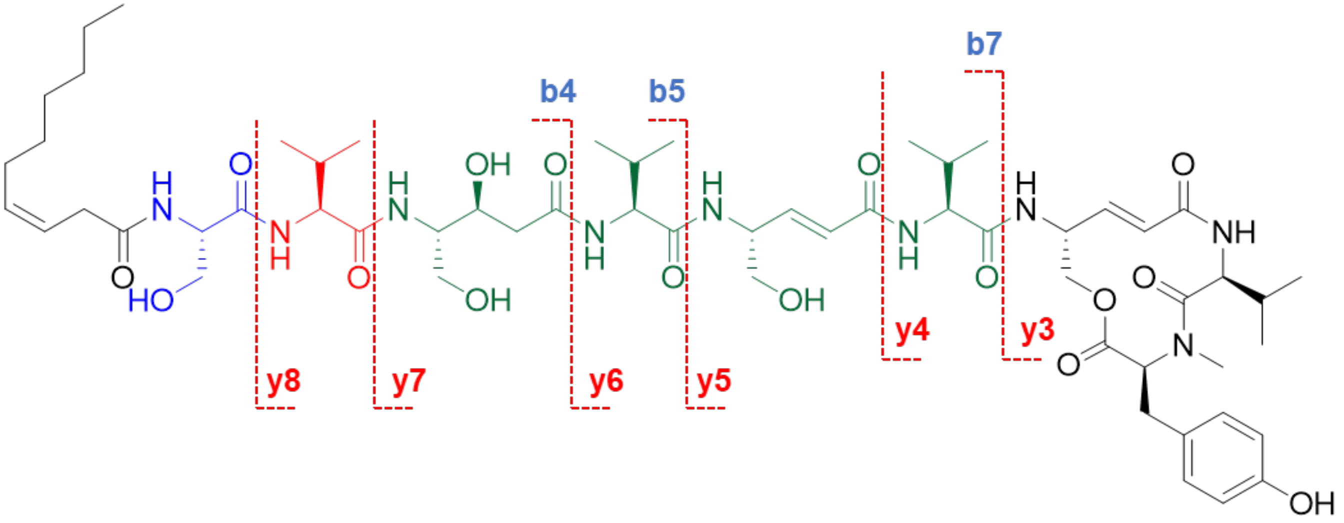


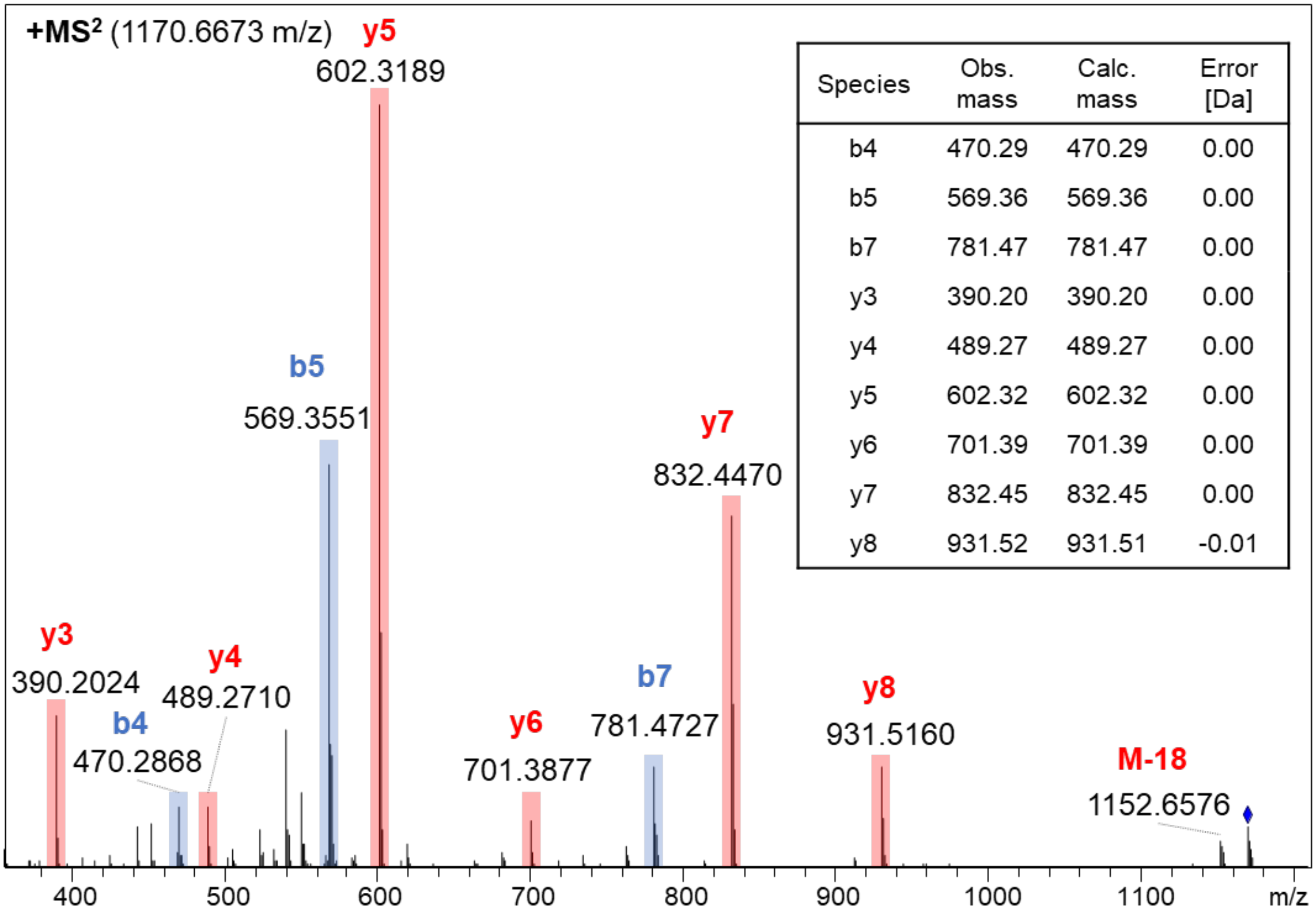


| Species | Obs. mass | Calc. mass | Error [Da] |
|---|---|---|---|
| b4 | 470.29 | 470.29 | 0.00 |
| b5 | 569.36 | 569.36 | 0.00 |
| b7 | 781.47 | 781.47 | 0.00 |
| y3 | 390.20 | 390.20 | 0.00 |
| y4 | 489.27 | 489.27 | 0.00 |
| y5 | 602.32 | 602.32 | 0.00 |
| y6 | 701.39 | 701.39 | 0.00 |
| y7 | 832.45 | 832.45 | 0.00 |
| y8 | 931.52 | 931.51 | -0.01 |

**Supplementary Fig. 20 | Characterization of thalassospiramide E7 (29).** MS$^n$ analysis.

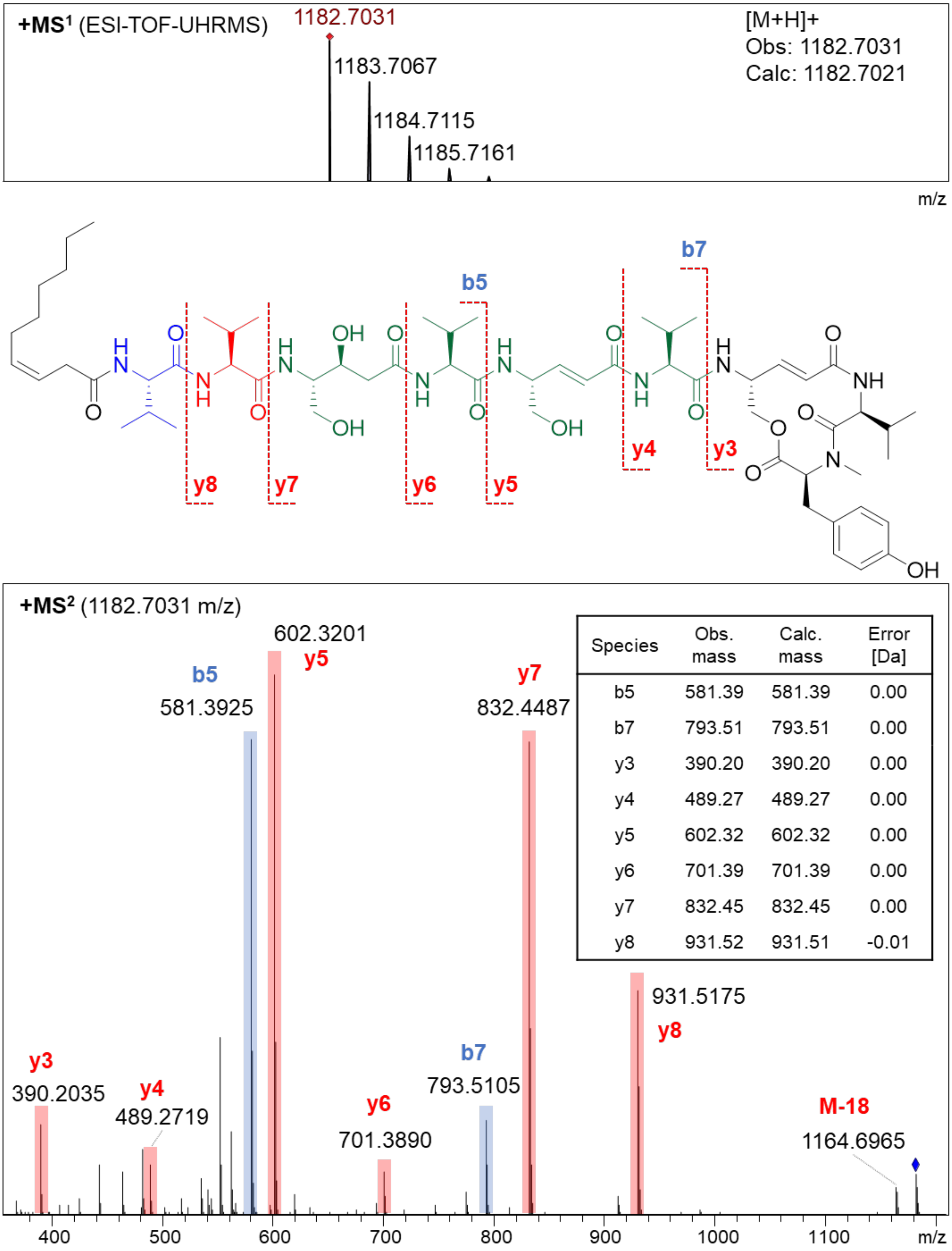


| Species | Obs. mass | Calc. mass | Error [Da] |
|---|---|---|---|
| b5 | 581.39 | 581.39 | 0.00 |
| b7 | 793.51 | 793.51 | 0.00 |
| y3 | 390.20 | 390.20 | 0.00 |
| y4 | 489.27 | 489.27 | 0.00 |
| y5 | 602.32 | 602.32 | 0.00 |
| y6 | 701.39 | 701.39 | 0.00 |
| y7 | 832.45 | 832.45 | 0.00 |
| y8 | 931.52 | 931.51 | -0.01 |

**Supplementary Fig. 21 l Characterization of thalassospiramide E8 (30).** $MS^n$ analysis.

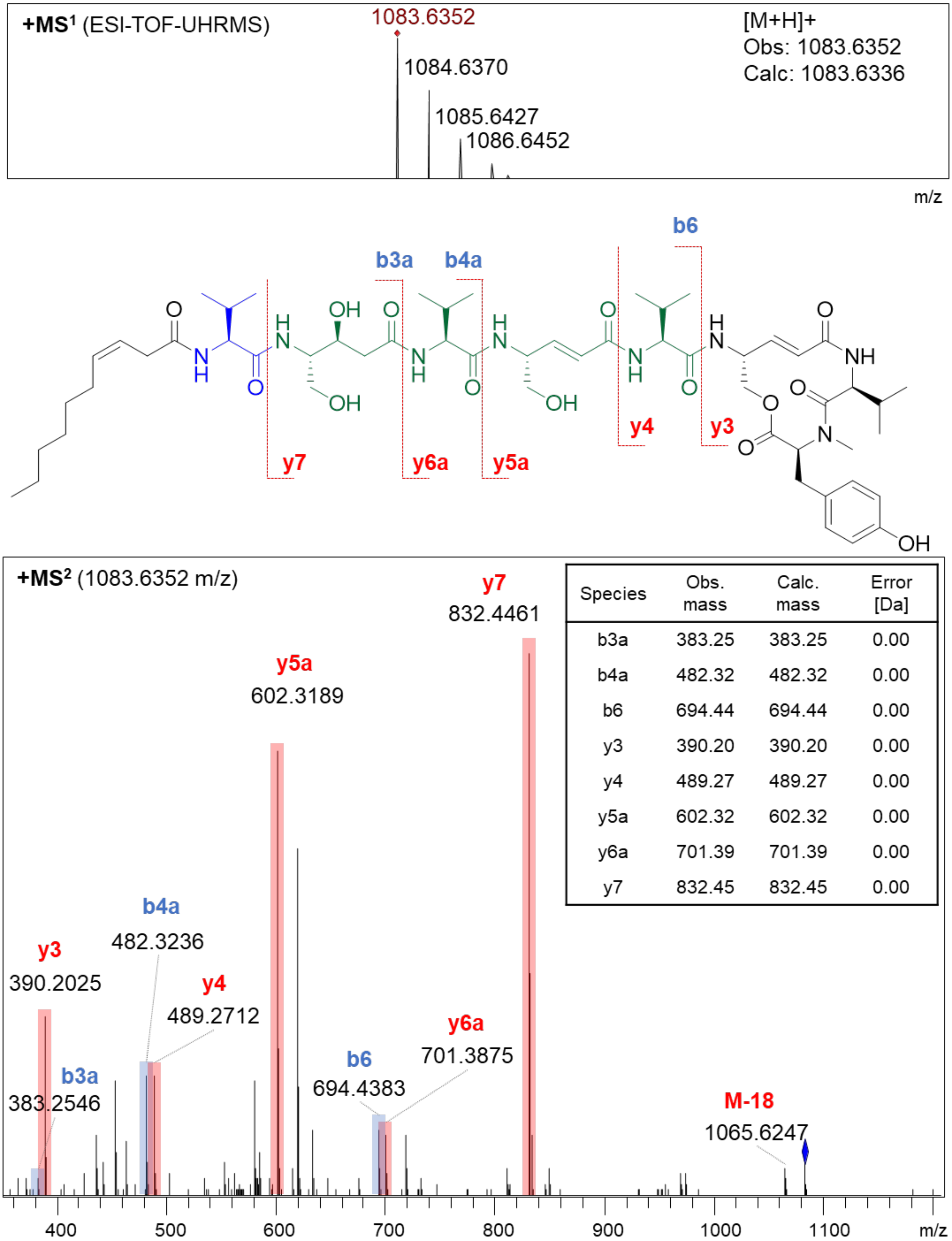


| Species | Obs. mass | Calc. mass | Error [Da] |
|---|---|---|---|
| b3a | 383.25 | 383.25 | 0.00 |
| b4a | 482.32 | 482.32 | 0.00 |
| b6 | 694.44 | 694.44 | 0.00 |
| y3 | 390.20 | 390.20 | 0.00 |
| y4 | 489.27 | 489.27 | 0.00 |
| y5a | 602.32 | 602.32 | 0.00 |
| y6a | 701.39 | 701.39 | 0.00 |
| y7 | 832.45 | 832.45 | 0.00 |

**Supplementary Fig. 22 | Characterization of thalassospiramide E9a (31).** MS$^n$ analysis. (NOTE: same spectra as E9b)

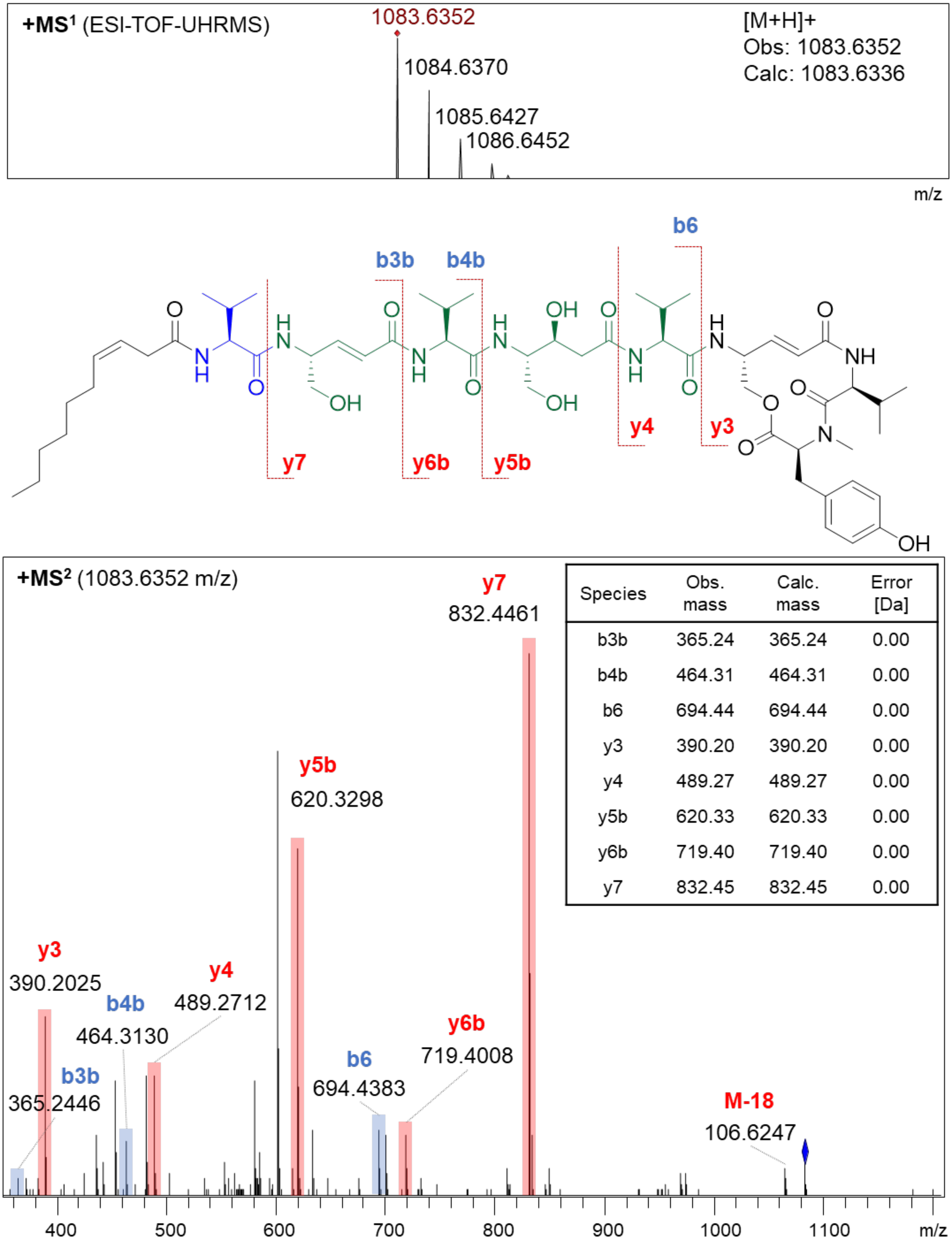


| Species | Obs. mass | Calc. mass | Error [Da] |
|---|---|---|---|
| b3b | 365.24 | 365.24 | 0.00 |
| b4b | 464.31 | 464.31 | 0.00 |
| b6 | 694.44 | 694.44 | 0.00 |
| y3 | 390.20 | 390.20 | 0.00 |
| y4 | 489.27 | 489.27 | 0.00 |
| y5b | 620.33 | 620.33 | 0.00 |
| y6b | 719.40 | 719.40 | 0.00 |
| y7 | 832.45 | 832.45 | 0.00 |

**Supplementary Fig. 23 | Characterization of thalassospiramide E9b (32).** MS$^n$ analysis.
(NOTE: same spectra as E9a)
